# Selective miniprotein inhibitors of Aurora-A kinase designed using interaction-motif scaffolding

**DOI:** 10.64898/2026.07.12.737516

**Authors:** Jennifer A. Miles, Bob Schiffrin, James Holder, Emily J. Wallis, Iain W. Manfield, Sean A. Burnap, Weston B. Struwe, Fanni Gergely, Richard Bayliss

## Abstract

Targeting kinase ATP-binding sites has produced many successful therapeutics, but selectivity remains a major challenge. We hypothesised that substrate-recognition surfaces could provide an alternative route to selective kinase inhibition. Here, we designed and tested structure-guided miniprotein inhibitors of Aurora-A using N-Myc as a template, a natural weak binder of the P+1 pocket. The most potent designs bind Aurora-A with single-digit nanomolar affinity, more than a thousand fold higher than the starting template, and selectively inhibit Aurora-A over Aurora-B in kinase assays and mitotic cells. Despite diverse architectures, successful binders targeting the substrate-recognition surface converged on a common strategy, coupling extensive engagement of the αG helix with activation loop stabilisation. Together these results show that selective kinase inhibition can be achieved by exploiting structurally well-defined active-state conformations rather than kinase-specific inactive states. Our work establishes a framework for converting weak substrate-recognition surface interactors into potent, selective kinase inhibitors through structure-guided protein design.

## INTRODUCTION

Protein kinases are master regulators of cellular signalling pathways and play central roles in human disease. Small molecule kinase inhibitors that target the ATP-binding site have transformed the treatment of kinase-driven cancers such as ALK+ non-small-cell lung cancer and chronic myeloid leukemia^1,2^. However, this strategy is limited to a single, highly conserved site, and achieving selectivity remains a barrier for many kinases. This challenge, and the relatively large chemistry effort required to overcome it, has limited the development of kinase inhibitors to a subset of targets with perceived high value. Thus, the landscape of kinase research and kinase inhibitors is skewed heavily towards Tyr kinases that are frequently mutated in cancer, and away from the diverse families of Ser/Thr kinases, many of which have poorly characterised functions^3^.

Although all protein kinases use ATP as a substrate, they are considerably more selective in their interactions with substrates and regulatory factors. These interactions are therefore attractive targets for selective kinase inhibitor development. However, the surfaces engaged in these interactions are shallow and extensive, which makes them challenging targets for small molecule inhibitors. There have been some notable successes, such as disruptors of the interaction between Aurora-A kinase and its activator TPX2^4,5^. There have also been positive developments in peptide-based inhibitors of protein kinases, but turning a weak native kinase-peptide interaction into a potent inhibitor requires complex design strategies and advanced peptide chemistry^5–7^. Miniproteins are well suited to this challenge because they can engage shallow and extended protein-protein interaction surfaces with high affinity and specificity^8^.

Recent advances in deep learning-based protein/peptide design have enabled the generation of miniproteins that bind to a designated target site, with nanomolar-picomolar affinities achievable without experimental optimisation in some cases^9–12^. Diffusion-based models such as RFdiffusion can design proteins completely *de novo*, or scaffold new proteins around one or more peptide motifs. Motif scaffolding-based approaches are especially suited to targeting protein-protein interactions, such as kinase-substrate interfaces, but remain underexplored. It is therefore unclear whether such strategies can be broadly employed to develop protein-based kinase inhibitors with high selectivity and cellular activity.

Aurora-A kinase is a Ser/Thr kinase with a well-established primary function in the regulation of mitosis. More recently, it has emerged as a key factor in resistance to cancer therapies, and as a regulator of the Myc family of transcription factors. Aurora-A has two closely related paralogs: Aurora-B which is also a ubiquitous mitotic regulator, and Aurora-C, which has restricted tissue expression. Most small molecule inhibitors of Aurora-A also inhibit Aurora-B because there is only one amino acid difference in their ATP sites that can be exploited for selectivity^13^. Aurora-A has therefore served as a model system to test alternative design strategies, exploiting regulatory and functional interactions with its binding partners such as TPX2, TACC3 and N-Myc^4,6,14–17^. While this effort has produced interesting and useful molecules, no single method has emerged that is both efficient and broadly applicable across kinases. Aurora-A is therefore an attractive model for testing protein-design-based strategies that target surfaces outside the ATP-binding site.

Here, we designed, built and tested motif-scaffolded miniprotein binders that directly inhibit Aurora-A by targeting its substrate-recognition surface. These designed binders exhibit high potency and selectivity for Aurora-A over Aurora-B *in vitro* and in cells. They also complement small molecule research tools by engaging structural features inaccessible to small molecules and by enabling applications such as selective isolation of Aurora-A complexes from cells. We envisage that this approach will be broadly applicable to the design of miniprotein inhibitors of protein kinases by targeting their underexplored substrate-binding surfaces.

## Results

### Design of motif-scaffolded Aurora-A binders

Although several structures of Aurora-A bound to natural protein activators and single domain antibody inhibitors have been resolved, there are no structures of this kinase with a peptide substrate at the active site^15,18–24^. However, the complex of Aurora-A kinase domain with a fragment of the N-Myc transactivation domain (TAD) shows an interaction with the P+1 pocket of the expected peptide substrate binding site on Aurora-A (Figure 1A, bottom right hand corner with N-Myc shown in blue)^22^. The region of N-Myc that is ordered in the structure (aa 61-89) was considered as a potential starting point for designing an inhibitor of Aurora-A activity. Residues 61-73 of N-Myc interact with the ⍺C helix and activation loop of Aurora-A, but this is not essential for the interaction. Residues 73-89 of N-Myc interact with the kinase P+1 pocket and form a helix that has been used as the basis for micromolar-affinity inhibitors of the N-Myc interaction, based on constrained peptides^16^. Therefore, this helix was selected as a starting point for inhibitory binder design.

**Figure 1.**
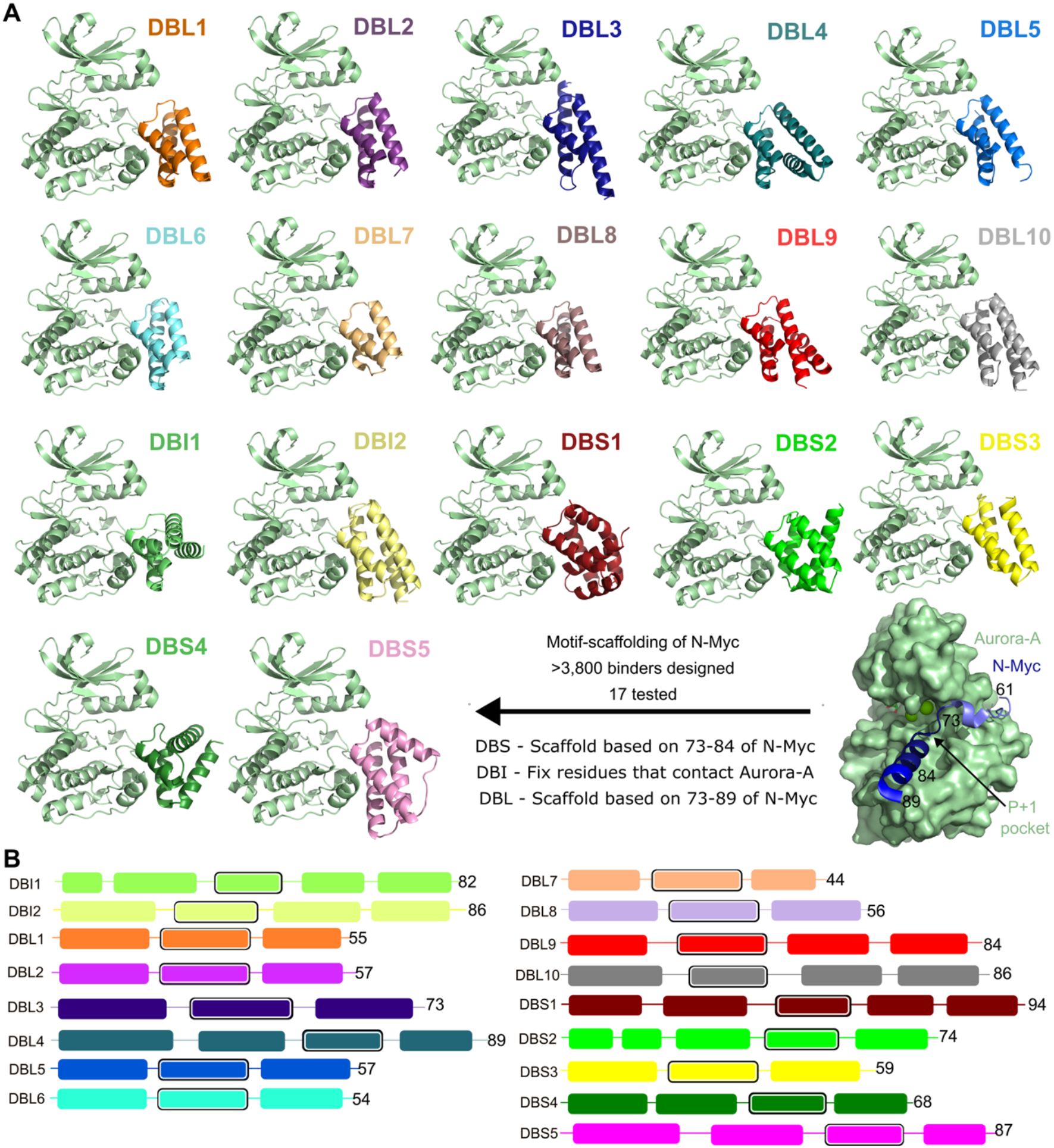
Design of Aurora-A miniprotein binders. **A:** AF2 predictions of binder:Aurora-A complexes. The binders are named based upon the region of the N-Myc sequence that has been used in scaffolding (DBS and DBL), or whether the residues in this region were retained in the final designs (DBI). The N-Myc:Aurora-A crystal structure (PDB: 5G1X) is shown in the bottom right-hand side, with the N-Myc region used for scaffolding highlighted (blue). **B:** Representation of the secondary structure of the 17 binders taken forward for *in vitro* analysis. The helix that was used as a template is highlighted with a black border.

RFdiffusion was used to design binders that scaffold either the core of the helical interface with Aurora-A (residues 73-84) or the full helix (residues 73-89). Between 10-50 residues were added to the N- and C-termini of this region to generate binders. 3,893 backbones were designed, with 1,967 and 1,926 backbones for residues 73-84 and 73-89, respectively. The hit rate was 2.5-3% based on an interface predicted aligned error (PAE) score of below 10^25^. 17 binder designs were prioritised based upon the compactness of the binder fold and AlphaFold2 (AF2) metrics of the predicted complex with Aurora-A: high mean binder pLDDT (>84) and low iPAE (<10) (Fig. 1A). The binders were named with relation to the region of N-Myc that was used in the original binder design (S, short – residues 73-84; L, long – residues 73-89) and the ‘L’ group was further split with designs in which the residues at the interface with Aurora-A were fixed at the ProteinMPNN sequence design stage (I, interface – residues Glu73, Pro75, Trp77, Val78, Thr79, Glu80, Glu84, Asn85, Trp88, and Gly89 of N-Myc). 2 of the 17 binders were in the I group, 5 were in the S group, and 10 were in the L group.

The AF2 models of all 17 binders place the scaffolded helix on the surface of Aurora-A, positioned to interact with the P+1 pocket (Figure 1A). The 17 designs were all-helical containing either 3 helices (8 designs), 4 helices (6 designs) or 5 helices (3 designs) (Figure 1A, B). The binders varied from 44-94 amino acids in length, and the scaffolded helix was always connected to at least one helix at both the N and C-terminus (Figure 1B).

### Motif-scaffolded binders potently inhibit Aurora-A

Genes encoding the 17 chosen binder sequences fused to an N-terminal His_6_-tag were synthesised, and the proteins overexpressed in *E. coli* and purified. 2 of the 17 binders expressed very poorly (DBL7 and DBL9), and so were not analysed further.

An ADP-Glo kinase assay was used to evaluate binder inhibition of Aurora-A kinase domain (122-403) activity. 8 of the 15 binders tested inhibited the activity of Aurora-A with sufficient potency so that an IC_50_ could be calculated (Figure 2A-C). The top two inhibitors were DBS1 and DBS5, with IC_50_ values of 12 nM and 16 nM, respectively (red and pink data in Figure 2B). Four of the DBL binders were also potent inhibitors, exhibiting IC_50_ values less than 100 nM (DBL2, DBL3, DBL5, DBL6). The two binders that had the interface residues fixed to match the original N-Myc sequence (DBI1 and DBI2) had the weakest effect on kinase activity. Neither exhibited inhibition of Aurora-A (yellow and green data in Figure 2A), suggesting that redesign of the interface residues of the scaffolded helix by ProteinMPNN was important for kinase inhibition.

**Figure 2.**
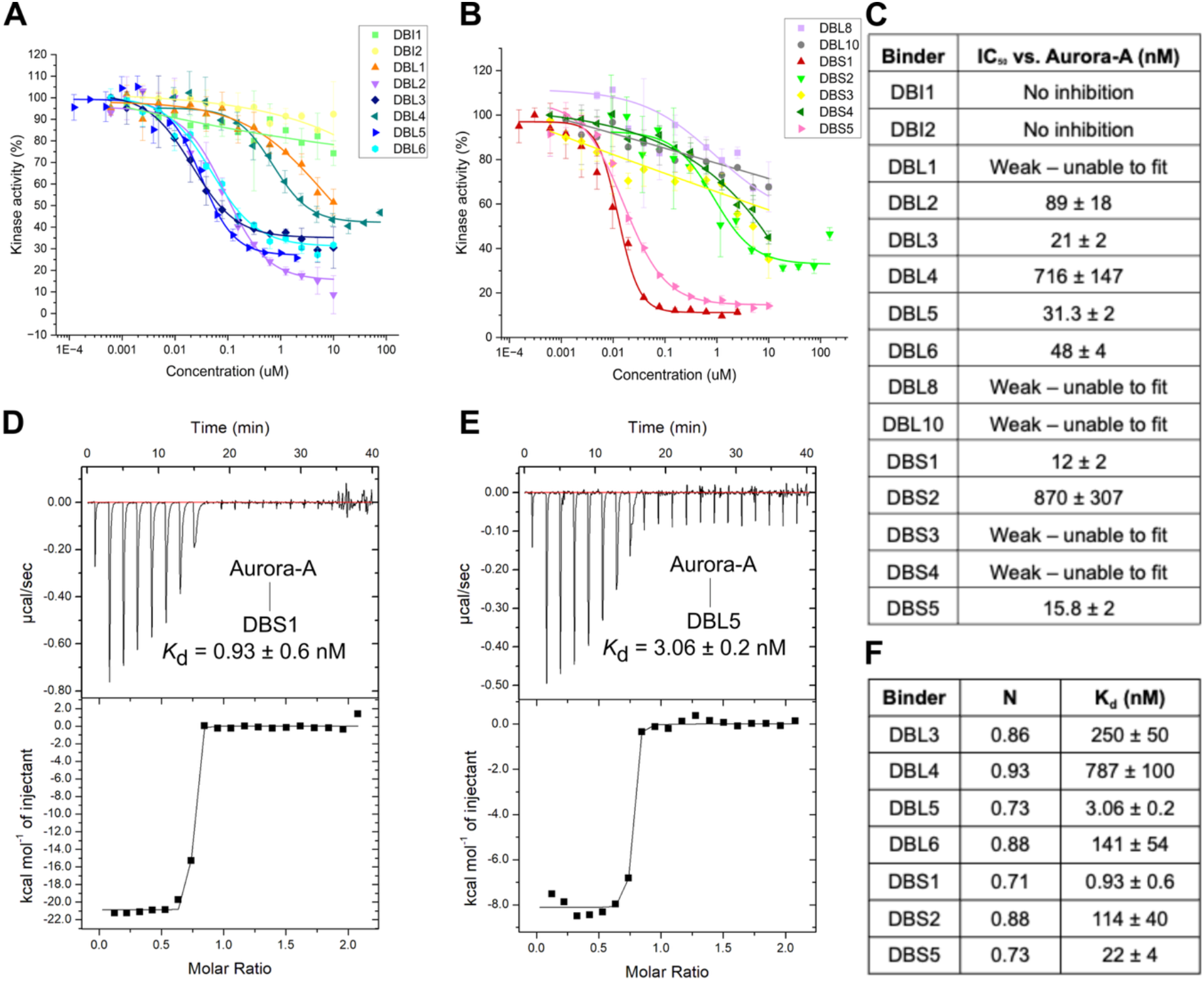
Inhibition and binding of Aurora-A by miniproteins. **A, B:** ADP-Glo assay to assess the effect of the designed binders on the ATPase activity of 50 nM Aurora-A WT 122-403, with 10 μM ATP and 100 μM kemptide as substrates. **C:** Summary table of IC_50_ values from A,B. **D-E:** Isothermal titration calorimetry experiment showing titration of Aurora-A kinase domain (122–403 C290A C393A D274N) into (D) DBS1, (E) DBL5. *K*_d_ was determined from two experimental repeats. **F:** Summary table of K_d_ values from ITC experiments.

Binding affinities for the interactions between the binders and unphosphorylated Aurora-A kinase domain were determined using isothermal titration calorimetry (ITC). Four of the binders produced weak signals that could not be fitted (DBL1, DBL2, DBL8 and DBS4). The affinities of the other binders for Aurora-A varied from 0.9 nM to 787 nM (Figure 2D-F, Figure S1). This is a ∼1000-fold increase in affinity when compared to the native interaction of N-Myc 61-89 binding to Aurora-A, which has a K_d_ of 12 μM when tested in a fluorescence polarisation-based assay^22^. Although absolute IC_50_ and K_d_ values are not strictly comparable, the trends across the binders are mostly consistent. DBS1 is the top-ranked by both values, it is the most potent inhibitor and tightest binder. Interestingly, while DBL5 and DBL3 have similar IC_50_ values (31 nM vs 21 nM), their K_d_ values are highly divergent (3 nM vs 200 nM).

### High-affinity binders extensively exploit the substrate-recognition surface

Next, we analysed the binder sequences and structures to investigate the gain in affinity of the binders compared to N-Myc, and the differences in affinity between them. To enable an accurate analysis of the binder/Aurora-A interactions, we set out to crystallize the complexes. To streamline the approach, we co-expressed the His-tagged binder with tag-free AurA^CAKD^. This is a kinase-dead, cysteine-free variant of the kinase domain that maximises protein yield while avoiding the risk of heterogeneous phosphorylation (AurA^CAKD^ : aa122-403 D274N C290A C393A)^26^. We determined crystal structures of nine of the binders in complex with AurA^CAKD^, from twelve attempted, at resolutions ranging from 1.79-2.9 Å (Figure 3A, Table S1). For most complexes, the difference between the AF2-modelled structure with the highest ipTM and the crystal structure was modest, with RMSDs across the binder atoms ranging from 0.5-1.3 Å, with the exception of DBL1 where the RMSD was 2.1 Å (Figure 3A).

**Figure 3.**
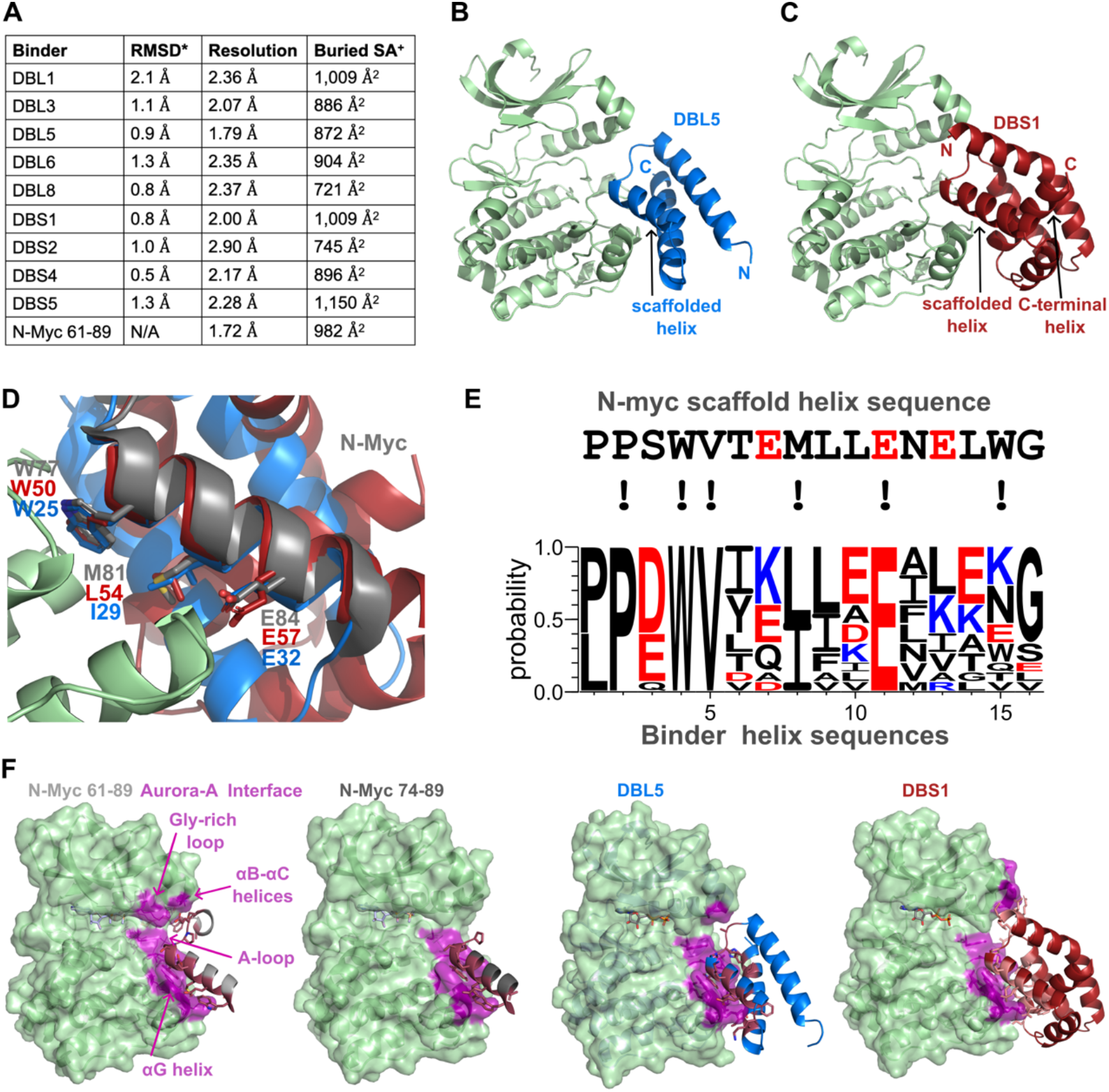
Sequence and structural properties of designed binders. **A:** Summary table of binder/ AurA^CAKD^ crystal structures. Root Mean Square Deviation (RMSD) compares the refined crystal structure model to an AF2 model, calculated using DALI pairwise. Buried surface area (SA) was calculated using PDBe PISA. **B:** Structure of DBL5 (blue) bound to the Aurora-A kinase domain (light green). **C:** Structure of DBS1 (dark red) bound to the Aurora-A kinase domain (light green). **D:** Comparison of two of the binder structures with the structures of N-Myc bound to Aurora-A, highlighting the critical interaction residues on the scaffolded helix. DBS1 is shown in dark red, with DBL5 in blue and N-Myc in grey. The structures have been aligned on the Aurora-A kinase domain, from the structure of N-Myc bound to Aurora-A (PDB: 5G1X, Aurora-A is shown in light green). **E:** Comparison of N-Myc template helix sequence with the sequences of the 17 designed binders. Critical interaction residues in N-Myc are indicated with “!”. **F:** The binding footprint of N-Myc and the binders on the surface of Aurora-A kinase domain (magenta).

We focussed initially on the crystal structures of the two highest affinity binders, DBS1 and DBL5. DBL5 is a simple 3-helix bundle in which the scaffolded helix is flanked at each end by one additional helix (Figure 3B). All three helices have a buried face and a solvent-exposed face. In contrast, DBS1 has a more complex topology, in which the scaffolded helix is flanked at each end by two helices, and its C-terminal helix is almost entirely buried (Figure 3C). The crystal structures are in close agreement with AF2 models (<1 Å RMSD, Figure S2B,C). The structures were compared with the original N-Myc bound structure that was used for scaffolding (PDB: 5G1X) (Figure 3D). Both DBS1 and DBL5 engage Aurora-A using a modified template N-Myc helix, with structurally equivalent residues mediating the interaction (Figure 3D). This confirms that, for these potent binders, the templated N-Myc helix is located at the core of the interface with Aurora-A, and that the design strategy was successful.

The binder sequences were aligned on the region corresponding to the N-Myc helix (Figure 3E, S2A). Most of the key residues that interact with Aurora-A were retained in all of the binders. Exceptions were the Met (at position 8 in Figure 3E), which was Leu or Ile in the binders, and the second Trp in the sequence (at position 15 in Figure 3E), which was more diversely substituted. In the context of the N-Myc/Aurora-A interaction, both Trp residues in the helix are critical for binding^22^, whereas in the binders only the first Trp was preserved. Indeed, only the two DBI group binders, where the second Trp was fixed, retained a bulky aromatic residue at this position. This Trp residue is clearly dispensable in the designed binders as neither DBI group binder substantially inhibited Aurora-A (Figure 2A). The second Trp in the N-Myc helix forms a H-bond with the side chain of Aurora-A Q335, a role that is preserved in the residues substituted in this position in DBS1 and DBS2. In contrast, the side chain of the residue at this position in DBL binders does not interact with Q335, but instead there are up to two H-bonds formed with the binder main chain nearby. However, there is no correlation between this feature and the potency of the binders. We conclude that sequence optimisation makes a modest contribution to the gain in affinity of the binders compared to N-Myc.

The helices that flank the scaffolded helix in the binders contribute in two key ways: they are required for the folding of the protein, and they expand the interaction surface. The scaffolded helix in the context of the N-Myc protein is disordered and therefore samples many different conformations, only a subset of which are competent to interact with Aurora-A^27^. In contrast, the additional helices that flank the scaffolded helix form extensive interactions with the face of the scaffolded helix that is solvent-exposed in N-Myc, which constrain it in the conformation competent to bind Aurora-A (Figure 3B,C). The additional helices in the binders also expand the surface of Aurora-A interaction beyond that of the scaffolded helix alone (Figure 3F, S2D). N-Myc 61-89 interacts with 22 residues from Aurora-A, using a 4 Å distance cut-off (Figure S2D). The regions of Aurora-A involved includes the Gly-rich loop, ⍺B and ⍺C helices, ⍺G helix and the activation loop. In contrast, the scaffolded N-Myc helix interacts with only 11 residues of Aurora-A, restricted to the C-lobe (activation loop and ⍺G helix). The binders interact with an expanded surface of Aurora-A, including with the N-lobe and more substantially with the activation loop (Figure 3F, S2D). However, the highest-affinity binder DBS1 has the same buried surface area as DBL1, a weak binder, and it is similar to the weakly-interacting N-Myc 61-89 (Figure 3F).

We examined the crystal structures carefully to identify features associated with differences in binding affinity. Four of the other crystal structures are of 3-helix bundles bound to Aurora-A: DBL1, DBL3, DBL6 and DBL8 (Figure S3A). In addition to the templated, central helix, all of the binders have a C-terminal helix that packs against the Aurora-A activation loop and ⍺G helix, and an N-terminal helix that does not interact with the kinase, but completes the fold of the binder. The C-terminal helix is of variable length, with the weak binders having short helices, and the strong binders having longer helices that engage the kinase ⍺G helix with additional interactions (Figure 4A). DBL5, the highest affinity binder of the DBL class, has an additional interaction with the activation loop, a hydrogen bond between His55 in DBL5 and the mainchain of Leu289 in the activation loop in Aurora-A (Figure 4B). In contrast, the C-terminal helix of DBL1 and DBL8 bends away from the surface of Aurora-A and their residues equivalent to His55 (e.g. Arg54 of DBL1) do not contact the activation loop (Figure 4B). To clarify the role of His55 in mediating a high affinity interaction with Aurora-A, a variant of DBL5 was generated in which alanine was substituted. As measured using the ADP-Glo assay, DBL5-H55A had an increased IC_50_ for Aurora-A of 425 nM, over 10-fold higher than that of DBL5 (Figure 4C). Notably, DBL3, which exhibits relatively weaker binding by ITC, does not form an equivalent interaction with the unphosphorylated activation loop (Figure S3B).

**Figure 4.**
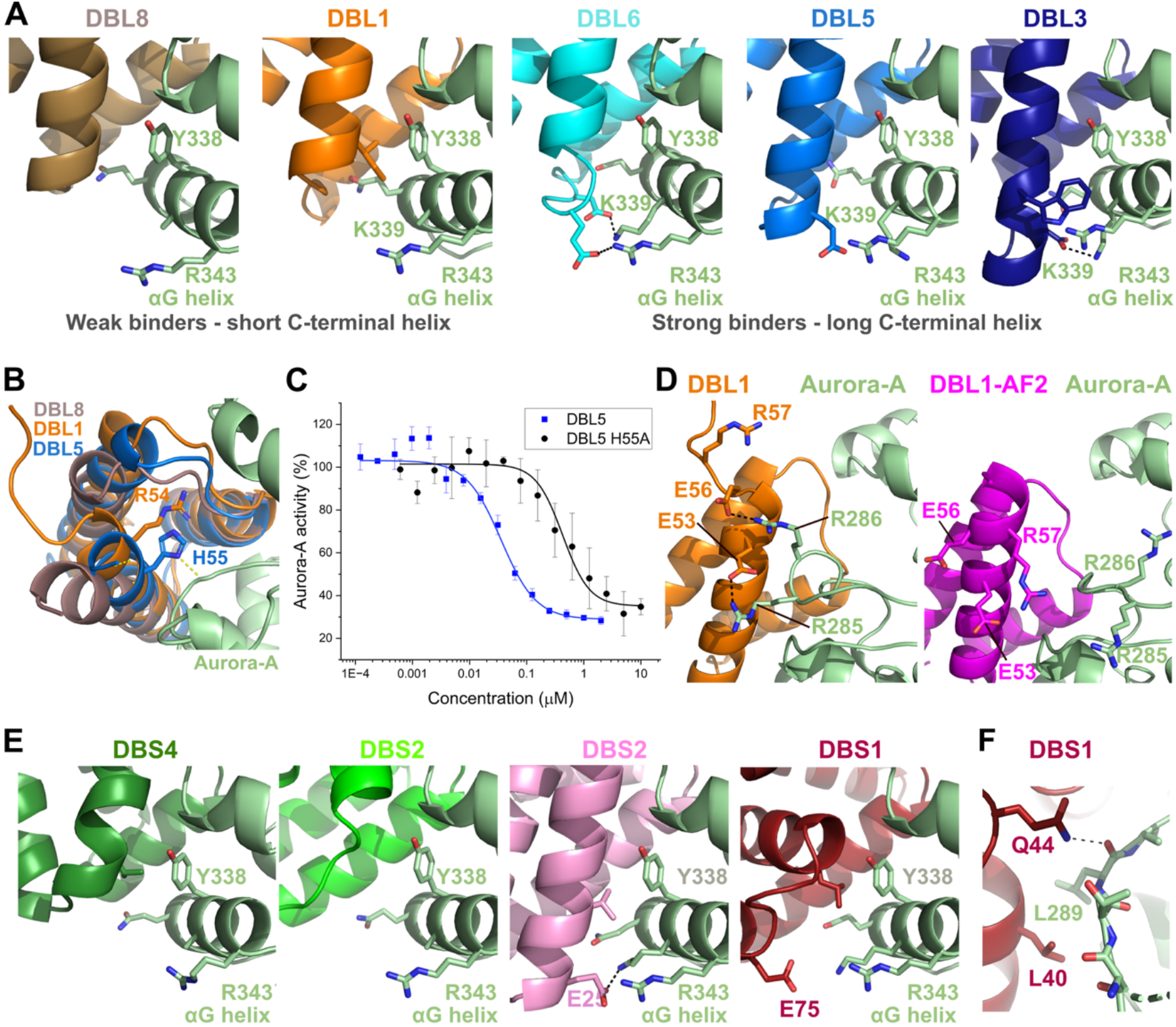
Relationship between miniprotein structure and binding affinity. **A:** Cartoon representation of the crystal structures of five DBL binders in complex with Aurora-A. The structures are aligned on the Aurora-A kinase domain (shown in pale green). **B:** Superposition of the crystal structures of three of the 3-helix binders bound to Aurora-A kinase. **C:** ADP-Glo assay to assess the effect of mutating DBL5 H55A on the activity of Aurora-A kinase. The data was fitted using Origin, with an IC_50_ of 425 nM ± 13 nM. **D:** (left) Magnified view of the crystal structure of DBL1/Aurora-A complex showing interactions with an unusual conformation of the activation loop. (right) Same view of an AF2 model of the DBL1/Aurora-A complex showing a predicted extended helix in DBL1. **E:** Cartoon representation of the crystal structures of four DBS binders in complex with Aurora-A. **F:** Magnified view showing the interaction of DBS1 Q44 with the main chain of the activation loop of Aurora-A.

The crystal structures also provide insights into the least successful binder designs because, at the very high concentrations used to form protein crystals, even weak binders can form complexes. Notably, the C-terminal helix of DBL1 is one turn shorter than in the model predicted by AF2 (Figure 4D). Two glutamic residues on the C-terminal helix of DBL1 interact with two arginine residues on the kinase activation loop, which adopts an unusual conformation. The incorrect modelling of this region of the binder is consistent with the relatively high RMSD between the crystal structure and the AF2 model (2.1 Å).

The other three structures comprised binders of 4 or more helices: DBS2, DBS4 and DBS5 (Figure S3C,D,E). Although these designs are somewhat more divergent than the DBL 3-helix binders, similar trends are apparent. The DBS binders with higher affinity for Aurora-A (DBS1, DBS5) have more extensive interactions with the ⍺G helix (Figure 4E). Analysis of the structures also suggest that higher potency could partially be mediated by interactions with the activation loop, which are missing from the weaker binders. In DBS1, the side chain of Gln44 forms a H-bond with the main chain of Leu289 (Figure 4F). There is also a potential H-bond between Thr10 in DBS5 and Thr288 in the activation loop of Aurora-A (Figure S3E). This suggests that interactions that stabilise the activation loop are favourable in the design of Aurora-A binders.

We conclude that the highest affinity binders exploit surface features of Aurora-A that are used by the natural binding partner N-Myc, such as the activation loop, but do so more effectively. They also extend the interface to establish new interactions with the αG helix. These features are combined with pre-ordering of the interacting residues to generate high affinity, folded binders from a low affinity, disordered protein.

### Designed miniproteins selectively target Aurora-A

Aurora-A is closely related to two other human kinases, Aurora-B and Aurora-C^28^. Aurora-C has only limited tissue expression, but most dividing cells express both Aurora-A and Aurora-B, which forms an obligate heterodimer with the chromosomal passenger protein INCENP^29^. Developing selective binders of Aurora-A has proven to be challenging because the kinase domains of these three proteins have similar sequences (Figure 5A)^13,28^.

**Figure 5.**
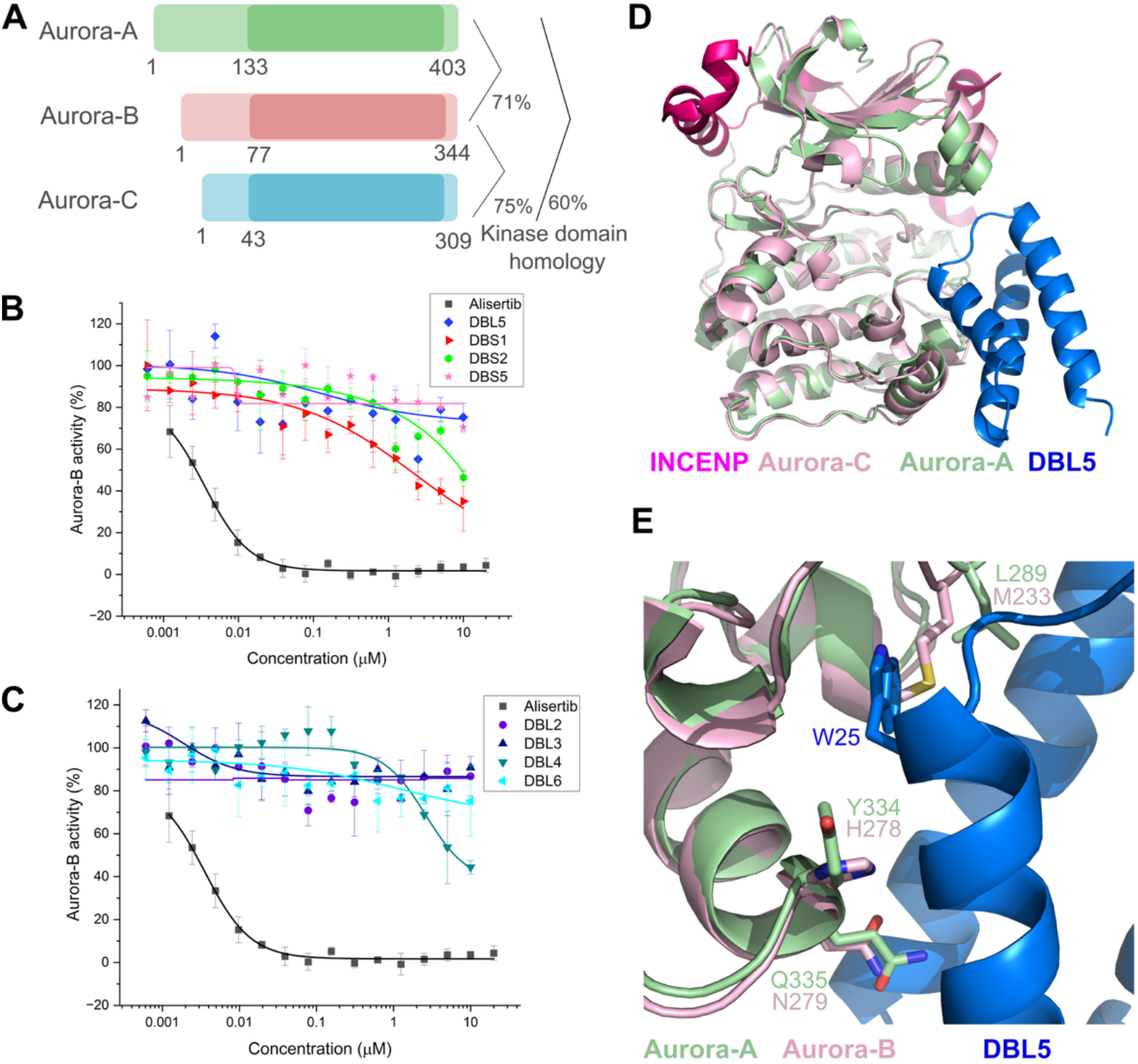
Assessment of miniprotein selectivity for Aurora-A over Aurora-B. **A:** Schematic visualisation of the sequence differences between Aurora-A, B and C. The kinase domain within each protein is highlighted in the darker region. The percentage identical amino acids between the kinase domains is indicated on the right-hand side. **B,C:** ADP-Glo assay to assess the effect of the designed binders on the ATPase activity of 150 nM Aurora-B in complex with INCENP, with 10 μM ATP and 100 μM kemptide substrates. Alisertib (20 µM) was used as a positive control for comparison. **D:** Superposition of crystal structures of Aurora-A/DBL5 and Aurora-C/INCENP (PDB code 6GR9). **E:** Magnified view of the superposition of the crystal structure of Aurora-A/DBL5 and an AF2 model of Aurora-B in an active conformation. Key residues at the interface with DBL5 that differ between Aurora family members are shown as sticks.

The designed binders were evaluated for Aurora-B inhibition in an ADP-Glo assay (Figure 5B,C). Human Aurora-B kinase domain (1-344) bound to INCENP (826-919) was incubated with binder ahead of the addition of ATP and the substrate peptide kemptide. The designed binders were very poor inhibitors of Aurora-B activity. The clearest Aurora-B inhibition was observed for DBS1, but this was at least 100-fold less potent than its inhibition of Aurora-A (Figure 5B, 2B). In this assay a control inhibitor, the small molecule ATP competitive compound Alisertib, was a potent inhibitor of Aurora-B.

Aurora-B is activated through binding of INCENP which itself becomes phosphorylated^29^. INCENP wraps around the N-lobe of the kinase domain and is not predicted to block the interaction of any of the designed binders with the Aurora-B/C kinase domain (Figure 5D). Using AF2 to generate models of the designed binders in complex with Aurora-B resulted in incorrect placement of the miniproteins and low ipTM scores. Closer examination of the binder interface with Aurora-A highlights differences at critical contact points (Figure 5E). In the P+1 pocket of the activation loop, Aurora-B/C have a methionine instead of a leucine (M233 vs L289), which would clash with the tryptophan that is preserved in all binders (e.g. W25 of DBL5). Furthermore, two of the contact points on the ⍺G helix, Y334 and Q335, are not conserved in Aurora-B/C. These differences are likely to be the major determinants of binder selectivity for Aurora-A over Aurora-B.

Next, we explored the functional consequence of the Aurora-A binders on mitotic cells (Figure 6). Standard biomarkers of Aurora kinase activity were selected: Aurora-A autophosphorylation on Thr288, and Aurora-B phosphorylation of Histone H3 Ser10. Small molecule Aurora-A and Aurora-B inhibitors were used for comparison. Overexpression of both DBL5 and DBS1 led to a significant decrease in Thr288 phosphorylated Aurora-A at the spindle poles and centrosomes (Figure 6A, B). In contrast, DBL5 had a negligible impact on Histone H3 Ser10 phosphorylation, while DBS1 reduced Histone H3 Ser10 phosphorylation levels, albeit its effect was significantly weaker than that of the Aurora-B inhibitor (Figure 6A,C).

**Figure 6.**
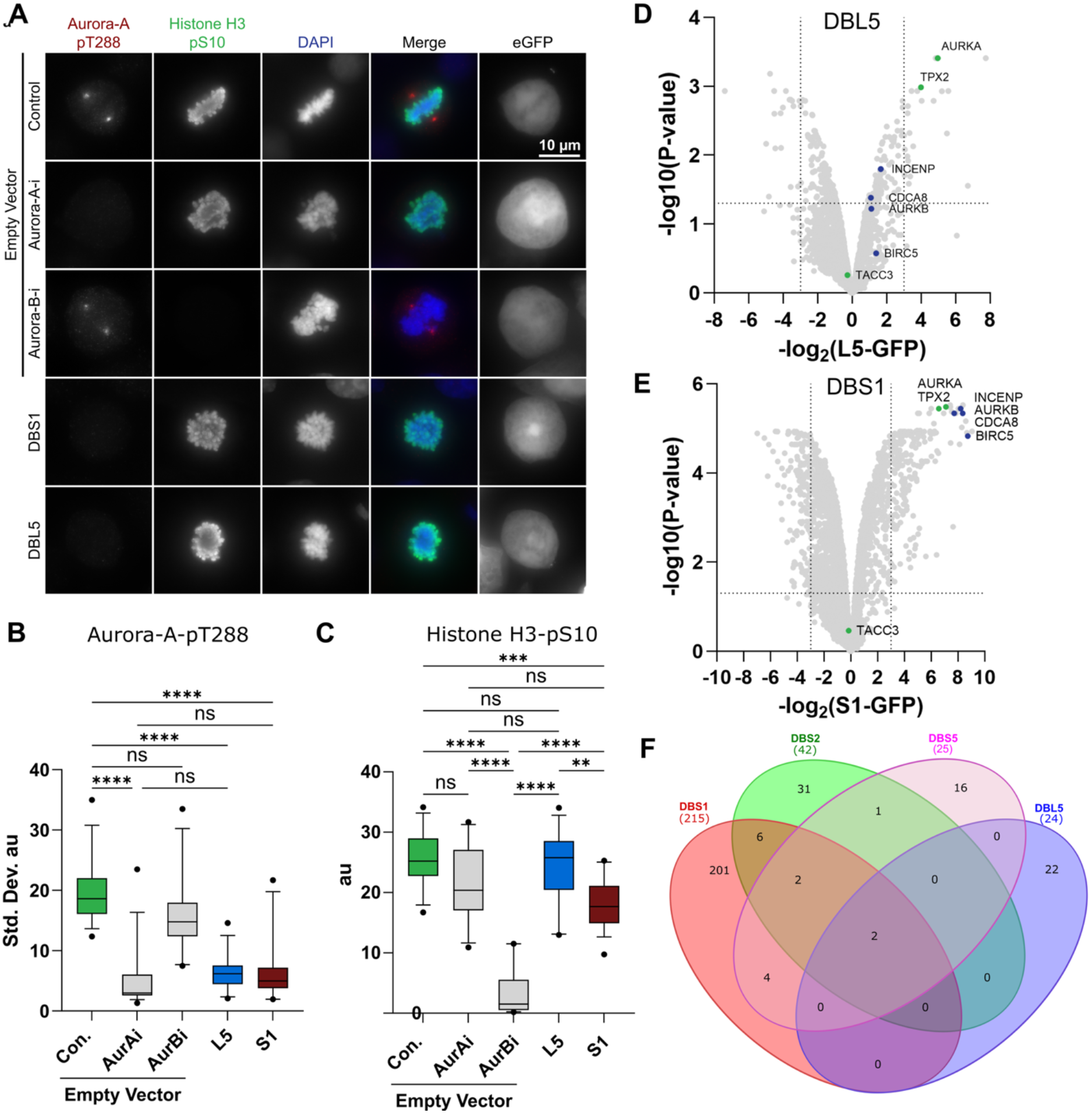
Miniproteins are selective inhibitors of Aurora-A in HeLa cells. **A**: Immunofluorescence of mitotic HeLa cells transfected with either GFP-empty vector or GFP-tagged miniproteins DBL5 and DBS1 48 h. Cells were treated with DMSO control, Aurora-A inhibitor (500 nM Alisertib) or Aurora-B inhibitor (2 μM ZM44743) for 25 min prior to fixation. Antibodies against Aurora-A-pT288 and Histone H3-pS10 are red and green in merged images, respectively, with DNA stained with DAPI (blue). **B-C** Box plots of (B) Aurora-A-pT288 spindle pole and (C) Histone H3-pS10 signal intensity in HeLa cells, measured in arbitrary units (au), with representative images shown in (A) (*n* = 3, ≥10 cells/biological replicate). **D-E** Volcano plot analysis showing log2 fold-change and significance (-log10 p-value) comparing protein enrichment with GFP-tagged designed binders (D) DBL5 or (E) DBS1 against GFP-only control. HeLa cells were transfected, as in (A), and 100 ng/ml nocodazole was added during the final 20 h of incubation to arrest cells in mitosis. GFP-immunoprecipitation was performed and samples analysed by LC-MS. **F:** Venn diagram summary of the numbers of proteins enriched following immunoprecipitation of 4 different GFP-tagged binders. The only proteins all four of the binders significantly enriched are Aurora-A and TPX2. Box plots in (B-C) indicate the median and interquartile ranges (25th–75th percentile) with coloured whiskers representing 5th–95th percentile ranges. *p* values are denoted as follows: \*\*\*\**p* < 0.0001, \*\*\**p* < 0.001, \*\**p* < 0.01, n.s not significant. Significance thresholds (D-F) were set at adjusted p-value <0.05 and log2 fold-change of >2.32 and are displayed as dotted lines.

To probe the specificity of the binders further, four of the GFP-tagged designs were selected for immunoprecipitation-mass spectrometry (IP-MS) analysis (DBL5, DBS1, DBS2, DBS5), using GFP as a control (Figure 6D, E and Figure S4A, B). DBS1 was noticeably more promiscuous than the others, with 215 protein interactors identified, whereas DBL5, DBS2 and DBS5 had only 24, 42 and 25 interactor proteins, respectively. All four of the binders interacted with both Aurora-A and TPX2, which were the only proteins in common between the four binders (Figure 6F). Consistent with this, a high-confidence AF3 model of the ternary complex between DBS1, TPX2 and Aurora-A (ipTM Aurora-A:TPX2 = 0.81; ipTM Aurora-A:DBS1 = 0.96) shows that DBS1 and TPX2 could bind Aurora-A simultaneously without steric clashes (Figure S4C). Furthermore, Aurora-A was the only kinase pulled out from the cells by DBS2, DBS5 and DBL5. DBS1 also interacted with Aurora-B, consistent with the weak inhibitory activity observed for Aurora-B in biochemical and cell-based assays. Two further kinases were identified as having an interaction with DBS1 – MARK2 (33% sequence identity) and MELK (30% sequence identity). The designed binder DBL5 was therefore confirmed to be a potent and selective binder of Aurora-A and its mitotic complex with TPX2, while DBS2 and DBS5 show a preference for this complex, and DBS1 is relatively less selective.

## Discussion

Here we show that the kinase substrate-recognition surface can be exploited to develop potent and highly selective inhibitors through computational protein design. Our goal was to establish and evaluate the approach using Aurora-A, which serves as a benchmark for the kinase inhibitor selectivity challenge because of its high similarity to Aurora-B^13^. Selectivity in kinase inhibitor development has often been established by targeting inactive states which are generally more structurally diverse than kinase active state conformations^30,31^. Here, we instead target the active state by engaging the substrate-recognition surface, which is only structurally well defined, and therefore amenable to rational design, in this conformation. Despite the structural similarity of active kinase conformations, we find that targeting this surface can nevertheless result in highly selective inhibitors. Indeed, three binders used for interactome proteomics showed selectivity for Aurora-A over Aurora-B, even though this was not an explicit design consideration. We rationalise that this selectivity arises because substrate-recognition surfaces have evolved to discriminate among diverse substrates and binding partners. Consequently, these regions exhibit substantially greater sequence and structural variation than the ATP-binding pocket. In human Aurora kinases, the ATP site has only three amino acid differences between Aurora-A and Aurora-B, only one of which is positioned to contribute directly to inhibitor selectivity^13^. In contrast, our structural data show that three Aurora-A residues are read by the binders to distinguish it from Aurora-B. Together, these findings establish the substrate-recognition surface as a previously underappreciated route to kinase selectivity. Whereas ATP-competitive inhibitors must distinguish between highly conserved nucleotide-binding pockets, designed binders can exploit the evolutionary diversification of substrate-recognition interfaces to achieve potent and selective inhibition^5,32^.

Although the highest-affinity binders were derived from distinct helical architectures, they converged on a common set of structural features that define a successful design strategy. Central to the enhancement of affinity is motif pre-organisation. Natural interactors of this surface, including kinase substrates and effectors such as N-Myc, engage Aurora-A through short motifs within disordered, conformationally heterogeneous regions^19,22,27^. By contrast, computational design embeds the motif within a stable folded domain, constraining it into a binding-competent conformation and reducing the entropic cost of binding. Beyond motif pre-organisation, the highest-affinity binders redistributed their contacts to engage the αG helix and activation loop in addition to the core substrate-recognition site. This expansion did not increase the overall buried surface area of the interaction, suggesting that the enhanced affinity reflects the quality and organisation of intermolecular contacts identified by exploration of the kinase surface rather than the formation of an overall larger interface. The recurrence of these interactions across distinct architectures suggests that they are key determinants of potent kinase inhibition.

Beyond demonstrating a route to potent kinase inhibition, this study also begins to define design rules for targeting substrate-recognition surfaces. Although emerging metrics such as ipSAE^33^ may help automate prioritisation of candidate binders, our results show that experimental validation remains essential and that predictive metrics should be complemented by design principles identified here, such as motif pre-organisation and extension of the interaction surface towards the αG helix and the active conformation of the activation loop.

Designed protein binders open up new ways to investigate and control protein kinases. While antibodies are the traditional tools that enable affinity purification and co-immunoprecipitation, their availability is limited to a restricted subset of proteins and epitopes. Here we show that designed binders can efficiently and selectively isolate Aurora-A from HeLa lysate. Efficient immunoprecipitation of Aurora-A from cell lysates is a known challenge that has previously been overcome by tagging the kinase with a protein for which well-established IP reagents are available, such as GFP^34^. A key advantage of miniprotein binders is that they enable the isolation of untagged Aurora-A. Moreover, the binder is selective for TPX2-bound Aurora-A complexes, consistent with its recognition of the active kinase conformation. Thus, the ability to recognise specific conformational states makes such binders precision tools for proteomics. We envisage using miniproteins to survey Aurora-A interactomes without the need for tagging, complementing or potentially replacing antibodies in selected applications.

Although intracellular delivery remains a challenge for many protein-based reagents, advances in protein and mRNA delivery have expanded the range of targets^35^ and enabled clinical translation of protein therapeutics such as the Myc-targeted miniprotein OMO-103^36^. The combination of potent inhibition, state-selective recognition and affinity capture demonstrated here establishes designed proteins as multifunctional reagents that can both interrogate and manipulate protein kinases. Crucially, this approach is also accessible and efficient. We generated potent and selective affinity reagents from an experimental screen of only 17 designs. Beyond Aurora-A, this approach may prove especially valuable for proteins that lack selective antibodies, affinity reagents or chemical probes, enabling the development of bespoke tools for previously inaccessible regions of the kinome.

## Methods

### Miniprotein design

Miniprotein binders to Aurora-A kinase were designed by motif scaffolding peptide sections of N-Myc. We used the high-resolution crystal structure of Aurora-A (residues 126-389) bound to a section of N-Myc (residues 61-89) (PDB ID: 5G1X, resolution 1.72 Å) as the target structure^22^. We selected either residues 73-84 or 73-89 of N-Myc as the motif to scaffold. Binder backbones were created with RFdiffusion^9^ keeping the N-Myc peptide fixed and extending the protein from both the N- and C-termini, stochastically sampling an extra 10-50 residues at both ends. Residues in contact with N-Myc were provided as hotspot residues. The whole chain of Aurora-A was provided as the target except for T288, which is phosphorylated in the 5G1X structure. An example RFdiffusion script is given below:

~~~
python ∼/RFdiffusion/scripts/run_inference.py \
“inference.output_prefix=output_files/aura_myc/bind” \
inference.input_pdb=auroa_myc_73-84.pdb \
’contigmap.contigs=[A127-287/A289-389/0 10-50/B73-84/10-50]’ \
‘ppi.hotspot_res=[A143,A172,A277,A296,A301,A333,A334,A335,A338]’ \
inference.num_designs=50
~~~

A total of 3,893 protein backbones were designed that scaffold N-Myc peptides (1,967 and 1,926 backbones for residues 73-84 and 73-89, respectively). Binder sequences were designed with ProteinMPNN and FastRelax using one relax cycle, and 1 or 2 sequences per backbone^37^. For binders that scaffold residues 73-89, an additional set of sequences were generated by fixing the identities of N-Myc residues that are in contact with Aurora-A in the crystal structure (residues 73, 75, 77, 78, 79, 80, 84, 85, 88, and 89). Structures of the Aurora-A/binder complexes were then predicted with AF2 using the af2_initial_guess protocol, which provides the designed structure as a template to the model. Filtering and selection of binders was performed by considering interface PAE (<10) and mean binder pLDDT (>84) scores, as well as visual inspection. All designs were all-helical, and we selected 17 binders containing 3 to 5 helices.

### Cloning

Codon optimised sequences for binders were ordered from GenScript including subcloning into pET28+ vector with a TEV-protease cleavable His-tag at the N-terminus. Q5-site directed mutagenesis (NEB) was used to produce the point mutant H55A into DBL5.

Designed binders were amplified with Q5-HF polymerase (M0491 NEB, primers ThermoFisher) and cloned into pJET1.2 vector according to manufacturer’s instructions. Binders were then digested and ligated (NEB) into pcDNA5-FRT/TO-eGFP vectors All constructs were N-terminally tagged except for DBS1 which was C-terminal. All constructs contained a 4x(GSSS) leaker between tag and protein coding sequence.

### Small scale expression of binders

The designed binders in pET28a+ were transformed into B834 RIL cells and the *E.coli* grown in 100 ml of LB at 37 °C. 0.5 mM IPTG was added to induce expression overnight at 20 °C. The cells were resuspended in 1 ml of lysis buffer (50 mM Tris-HCl pH 7.5, 500 mM NaCl, 20 mM Imidazole, 1 mM TCEP) and protease inhibitor tablets (Roche). Cells were lysed with sonication and the soluble fraction collected at 13,000 rpm for 10 min. The soluble fraction was bound to a His SpinTrap (Cytiva) equilibrated in lysis buffer. The spin column was washed with lysis buffer before the bound protein was eluted in 200 μl of lysis buffer with 300 mM Imidazole. The imidazole was removed with dialysis into 50 mM Tris-HCl pH 7.5, 250 mM NaCl, 0.5 mM TCEP. The purity of the binders was assessed using SDS-PAGE, followed by measurement of the concentration on the NanoDrop (Thermo Fisher Scientific). The binders were flash frozen and stored at −80°C.

### Large scale expression of binders

The designed binders in pET28a+ were transformed into B834 RIL cells and the *E.coli* grown in 2 litres of LB at 37 °C. 0.5 mM IPTG was added to induce expression overnight at 20 °C. The cells were resuspended in 10ml per litre of lysis buffer (50 mM Tris-HCl pH 7.5, 500 mM NaCl, 20 mM imidazole, 1 mM TCEP) and protease inhibitor tablets (Roche). Cells were lysed with sonication and the soluble fraction collected at 17,000 rpm for 45 min. The soluble was incubated with 3 ml of Ni-NTA resin (Qiagen) equilibrated in lysis buffer for 1 h. The resin was washed in 20 CV of lysis buffer before elution of binder in 4CV of lysis buffer with 300 mM Imidazole. The his-tag was removed by overnight dialysis into 50 mM Tris-HCl pH 7.5, 200 mM NaCl, 0.5 mM TCEP in the presence of TEV protease. The dialysed protein was incubated with 3 ml of Ni-NTA equilibrated in dialysis buffer to remove the uncut protein. The flow through and a wash containing tag free protein was concentrated in a 5 kDa cut off concentrator (Amicon) and loaded onto the SD200 equilibrated in 50 mM Tris-HCl pH 7.5, 200 mM NaCl, 0.5 mM TCEP. Fractions containing the binder were concentrated, flash-frozen fand stored at −80 °C.

### Co-purification of miniprotein:Aurora-A complexes

The designed binders in pET28a+ and Aurora-A D274N C290A C393A 122-403 in pCDF were co-transformed into B834 RIL cells. The proteins were co-expressed in 4 litres of LB at 37 °C and expression induced overnight with 0.5 mM IPTG at 20 °C. The cells were resuspended in 10 ml (per litre of culture) of lysis buffer (50 mM Tris-HCl pH 7.5, 250 mM NaCl, 20 mM Imidazole, 5 mM MgCl_2_, 10% glycerol) and protease inhibitor tablets (Roche). Cells were lysed with sonication and the soluble fraction collected at 17,000 rpm for 45 min. The soluble fraction was filtered through 0.45 µm filters before loading onto a HisTrap HP column (Cytivia) equilibrated in lysis buffer. This was attached to the akta pure and washed with 10 CV of lysis buffer before elution of anything bound in a gradient of lysis buffer containing 500 mM Imidazole. The His-tag was removed by overnight dialysis into 50 mM Tris-HCl pH 7.5, 250 mM NaCl, 10% glycerol, 5 mM MgCl_2_, 0.5 mM TCEP, in the presence of 1mg of TEV protease. The cleaved protein was rebound to the HisTrap HP equilibrated in dialysis buffer. The complex was eluted using a gradient of 0.5M imidazole as the cleaved Aurora-A still somewhat interacts with Ni-NTA. The complex was concentrated using a 30 kDa MWCO concentrator and loaded onto an SD200 column equilibrated in 50 mM Tris-HCl pH 7.5, 250 mM NaCl, 10% glycerol, 0.5 mM TCEP, 5 mM MgCl_2_. Fractions containing the complex of binder and Aurora-A were concentrated using a 30 kDa MWCO concentrator, flash frozen and stored at −80 °C.

### Production of Aurora-A for ITC and ADP-Glo

Aurora-A 122-403 for use in ADP-Glo assays was co-expressed with lambda phosphatase and purified as detailed previously^20^. Aurora-A D274N C290A C393A 122-403 for use in ITC and crystallisation was overexpressed and purified from *E.coli* as detailed previously^20^.

### Crystallography

ADP was added to the binder:Aurora-A complexes (15-20 mg/ml) to a final concentration of 5 mM and the samples left on ice for 1 h before crystallisation trials were set up. The complexes were screened against a range of commercial crystallization matrices. Drops were set at 1:1, 1:1.5 and 1.5:1 ratios of complex:precipitant in MRC sitting drop plates using a Mosquito LCP crystallization robot (STP Labtech) and incubated at 18 °C.

Crystals grew of 9 complexes in various conditions. Crystals were flash-frozen in liquid nitrogen. Diffraction data were collected from a single crystal at Diamond Light Source (Oxford, UK) on beamlines i03 and i04. Autoprocessed data from xia2 “3-daii” pipeline at Diamond Light Source were used for structure determination (Table S1)^38^. Molecular replacement was performed in PHASER using the AF2 model of the complex of Aurora-A and the designed binder^39,40^. Clear difference density was observed for the designed binder and this was modelled in using Coot^41^. Subsequent rounds of iterative refinement were performed using Phenix^42^ and Coot. MolProbity^43^ and PDB-Redo^44^ were used to determine structure quality. Key contributions to the interaction between Aurora-A and designed binder were identified by analysis of the structure using PDBePISA (Proteins, Interfaces, Structures and Assemblies)^45^. Root Mean Square Deviation (RMSD) was calculated using DALI pairwise^46^.

### ITC

Binding affinities were determined using an iTC200 microcalorimeter (MicroCal from Malvern Panalytical). Designed binders and Aurora-A 122-403 C290A C393A D274N were dialysed overnight into 20 mM Tris-HCl pH 7.5, 200 mM NaCl, 10% glycerol, 5 mM MgCl_2_. Experiments were carried out at 25 °C. The syringe contained 70 μl of 250 μM Aurora-A 122-403 (D274N C290A C393A), and the cell contained 205 μl of 25 μM designed binder. 2 μl injections were applied every 180 sec. Microcal Origin software version 7.0 was used to determine the dissociation constants (*K*_d_) and n values. All measurements were repeated at least twice.

### ADP-Glo

An ADP-Glo assay kit (Promega) was used to test the effect of the designed binders on the ATPase activity of Aurora-A kinase or Aurora-B kinase. Purified and tag-free designed binders were titrated into 50 nM Aurora-A 122-403 dephosphorylated WT protein or 150 nM Aurora-B 1-344 bound to INCENP 826-919 (Dundee reagents DU4009,DU930) in a buffer of 40 mM Tris-HCl pH 7.5, 150 mM NaCl, 10 mM MgCl_2_, 1 mM DTT, 0.1 mg/ml BSA, 0.01% Tween 20. Reactions were initiated by the addition of 10 μM ATP and 100 μM kemptide as substrate (Cambridge Bioscience Ltd). The reactions were incubated at room temperature for 60 min before 5 μl was transferred into 5 μl of ADP-Glo Reagent (Promega) in a 384-well low volume low binding plate (Greiner Bio-one). After a further 40 min at room temperature, 10 μl of kinase detection reagent was added to each well and the reaction allowed to incubate for an additional 30 min. Luminescence was measured on a HIDEX Sense plate reader (HIDEX) using a 1 sec integration time. Control data was subtracted from the luminescence and three repeats were averaged before being fitted using Origin (OriginLab), where IC_50_ values were calculated using the Levenberg-Marquardt Algorithm on a nonlinear curve fit.

### AF2 and AF3 modelling

Two seeds were used on AF2 with maximum 24 recycles to generate the models of Aurora-A 122-403 or Aurora-B in complex with the designed binders.

### Immunoprecipitation

2 × 10^6^ HeLa cells were seeded in a 15 cm dish and incubated at 37 °C, 5% CO_2_ for 48 hr. Cells were then transfected as follows (per dish): 1 µg pcDNA5-FRT/TO-eGFP-designed binder or empty vector constructs with 7 µg pBluescript empty vector construct + 24 µl Lipofectamine 3000 (Invitrogen) and 1 ml Opti-MEM, according to manufacturer’s instructions. Transfected cells were incubated for 48 hr, with 100 ng/ml nocodazole (CalBiochem, 487928) added for the final 20 hr. Cells were collected by mitotic shake-off and washed twice with PBS, once with Opti-MEM, and then incubated in fresh Opti-MEM for 30 min to rebuild mitotic spindles. For prior washing steps all spins were at room temperature, 5 min, 400*g* and all buffers were pre-equilibrated to 37 °C, 5% CO_2_.

Cells were pelleted lysed on ice for 30 min in the following buffer: 20 mM Tris-HCl (pH 7.5), 150 mM NaCl, 0.1 % NP-40, 0.1% Sodium Deoxycholate, 40 mM β-glycerophosphate, 10 mM NaF, 0.3 mM Na_3_VO_4_, 2 mM MgCl_2_, Benzonase (1 μl/ 5 ml), 100 nM okadaic acid, protease inhibitor cocktail (1:300, Merck P8340) and phosphatase inhibitor cocktail 3 (1:200, P0044). Samples were clarified (15 min, 14,000 g, 4 °C), the supernatant collected and protein concentrations measured by Bradford assay.

For each sample, 2.8 mg of clarified lysate was taken and the volume adjusted to 300 μl before a 10 μl input sample was taken. ChromoTek GFP-Trap Magnetic Agarose beads (Chromotek, gtma, 15 μl bead slurry/sample) were washed three times in lysis buffer and resuspended in 200 μl of lysis buffer per sample. Beads were added to the normalised lysates giving a final volume of 500 μl and incubated, while rotating, for 2 h at 4°C. The beads were washed three times with lysis buffer before either preparing for mass spectrometry analysis. All bead washes were done using a magnetic rack on ice.

### Immunofluorescence microscopy

100,000 HeLa cells were seeded in 6 well plates and grown 24 h and transfected as follows: 200 ng pcDNA5-FRT/TO-eGFP-designed binder construct, with 600 ng pBluescript empty vector construct, 3 µl Lipofectamine3000 (Invitrogen) and 200 μl Opti-MEM. Transfected cells were incubated for 48 hr prior to fixation in 4% paraformaldehyde for 12 min at room temperature. Coverslips were then quenched in 50 mM NH_4_Cl (10min), washed twice in PBS and further incubated in pre-cooled methanol for 5 min at 20°C. Prior to staining all coverslips were permeabilised in 0.2 % (v/v) Triton X-100 in PBS for 5 min and then blocked in 2% BSA, 0.1% (v/v) Tween in PBS. Blocking and all antibody incubation steps were carried out for 1 h unless otherwise stated. Primary and secondary antibodies (Life technologies) were diluted 0.1% Tween, PBS, with secondary staining being conducted in the dark. Coverslips were gently washed three times in 0.2% Tween, PBS between antibody incubation steps. 500 ng/ml DAPI (Sigma, D9542) was included during the secondary antibody incubation to stain DNA. Coverslips were mounted on glass slides (SuperFrost Ultra Plus, Thermo Scientific) with Prolong Diamond antifade mountant (Invitrogen), allowed to cure overnight at room temperature and stored at 4°C. A standard upright microscope system (BX61, Olympus) was used to image coverslips using GFP/Alexa Fluor 488, Cy3/Alexa Fluor 555, Cy5/Alexa Fluor 647 and DAPI filter sets (Chroma Technology Corp.). Images were taken and Z-projected using MetaMorph 7.5 imaging software (Molecular Devices) and a 2048 × 2048-pixel complementary metal oxide semiconductor camera (PrimΣ, Photometrics). Illumination was provided by an LED light source (pE300, CoolLED Illumination Systems). Z-stacks were obtained at a spacing of 0.4 µM throughout the volume of the cell.

### Mass spectrometry sample preparation and analysis

On-bead tryptic digest was performed as follows. Beads were washed twice in PBS, then twice in 100 mM ammonium bicarbonate (AMBIC) pH 8.0. Samples were digested with 100 ng trypsin (ProMega, V5280), in 100 mM AMBIC, for 4 h prior to the addition of fresh trypsin for an overnight incubation. Digestion steps were performed at 37 °C. Reactions were quenched with 5 % formic acid (v/v) to a final concentration of 0.5 %.

Peptides were desalted on homemade C18 stage tips (Empore, 66883-U). Columns were washed twice in 100 μl of 100 % acetonitrile (ACN) (v/v), and equilibrated with two washes with 100 μl of 0.1 % trifluoroacetic acid (TFA) (v/v). All spin steps were performed at 3000*g*, 2 min, unless otherwise stated. Sample was loaded onto the column (1500g, 5 min), the flow-through was reloaded onto the column twice. Columns were washed three times with 100 µl 0.1 % TFA and peptides eluted into fresh tubes with 60 µl 50 % ACN, 0.1 % TFA (v/v). The elution step was repeated once. Peptides were dried under vacuum and resuspended for mass spectrometry analysis.

Peptide samples were analysed using a NanoElute 2 UHPLC coupled to a timsTOF Ultra 2 mass spectrometer using a CaptiveSpray ionisation source (Bruker). Peptides were separated on a 75 μm × 25 cm IonOptiks Aurora Ultimate C18 analytical column (IonOptiks). Buffer A was 0.1% FA in H2O and buffer B was 0.1% FA in LC-MS grade acetonitrile (MeCN). A 30 or 120-minute linear gradient (0% to 40% buffer B) was used for elution.

Source parameters were set to a capillary voltage of 1,600 V, a dry gas flow of 3.0 L/min, and a dry gas temperature of 200 °C. Data was acquired in dia-PASEF mode over an m/z range of 400–1,000 and an ion mobility range of 1/K₀ = 0.64–1.45 Vs/cm2. The TIMS analyzer was operated at a 100% duty cycle with 100 ms accumulation and 100 ms ramp times (acquisition cycle time = 0.96 s). Collision energies were linearly stepped from 20 to 59 eV as a function of increasing ion mobility.

DIA data were quantified using DIA-NN (v2.3.0)1 with a spectral library predicted from FASTA (Homo sapiens, UP000005640_9606)^47^. Mass accuracy settings for MS1 and MS2, as well as the scan window, were determined automatically by DIA-NN. Precursor and protein group identifications were filtered at 1% FDR. Match between runs and protein inference were enabled. QuantUMS (precision) was used for quantification with retention time dependent cross run normalisation. Quantification matrices for precursors, protein groups, and gene groups were exported with 1% FDR filtering. N-terminal methionine excision and carbamidomethylation of cysteines were set as fixed modifications. The protein group matrix report was taken forward for statistical analysis.

### Statistical analysis of proteomic data

Mass spectrometry intensity data were analysed in R version 4.4.2. Protein intensity values and gene names were imported from the raw proteomic data. Data was filtered, only retaining proteins for downstream analysis if they were detected in at least two replicates in either experimental group, with ≥3 peptides identified across all conditions. Remaining intensity values were log2-transformed and median-centred on a per-sample basis. Missing values were imputed randomly from a normal distribution downshifted from the observed sample mean by 2.32 standard deviations with a width of 0.3 times standard deviation. A fixed random seed was used to ensure reproducibility.

Statistical analysis was performed using the limma package. A linear model was fitted to the imputed, median-aligned log2 intensities, comparing immunoprecipitations for GFP-tagged designed binder vs GFP-empty vector control using empirical Bayes moderation. Protein abundances were considered significantly different if they had an adjusted P value <0.05 and an absolute log2 fold change ≥3. Statistical analysis was performed on the whole dataset, while the volcano plots show only known centrosomal proteins for clarity.

### Sequence analysis

The binder sequences were aligned using MAFFT^48^ and a WebLogo^49^ was generated covering the region of the binders that was used for the original scaffolding. The sequences of the human Aurora-A kinase domain (Uniprot O14965 122-403), Aurora-B kinase domain (Uniprot Q96GD4 77-327) and Aurora-C kinase domain (Uniprot Q9UQB9 40-293) were aligned using MAFFT.

## Acknowledgements

This research was supported by the Biotechnology and Biological Sciences Research Council (SPIDR: BB/V003577/1, BB/V003577/2 for RB) and Medical Research Council (UKRI585 for FG). B.S. acknowledges support and funding from the Astbury Centre for Structural Molecular Biology. We acknowledge Diamond Light Source for time on beamlines i03 and i04 under proposal mx35120.

## Author contributions

J.A.M., B.S, F.G and R.B. designed the study; J.A.M. and B.S. carried out protein design; J.A.M. performed protein biochemistry and crystallography (with support from E.J.W) and analysis. J.H. performed cell biology, mass spectrometry sample preparation and data analysis; I.W.M. carried out isothermal titration calorimetry; S.A.B. and W.B.S. carried out proteomics. J.A.M., J.H., F.G. and R.B, wrote the manuscript. All authors have reviewed and approved the manuscript.

## References

1. Elshatlawy, M., Sampson, J., Clarke, K., and Bayliss, R. (2023). EML4-ALK biology and drug resistance in non-small cell lung cancer: a new phase of discoveries. Mol Oncol 17, 950–963. 10.1002/1878-0261.13446.

2. Guilhot, F., and Hehlmann, R. (2025). Long-term outcomes of tyrosine kinase inhibitors in chronic myeloid leukemia. Blood 145, 910–920. 10.1182/blood.2024026311.

3. Gomez, S.M., Axtman, A.D., Willson, T.M., Major, M.B., Townsend, R.R., Sorger, P.K., and Johnson, G.L. (2024). Illuminating function of the understudied druggable kinome. Drug Discov Today 29, 103881. 10.1016/j.drudis.2024.103881.

4. Stockwell, S.R., Scott, D.E., Fischer, G., Guarino, E., Rooney, T.P.C., Feng, T.S., Moschetti, T., Srinivasan, R., Alza, E., Asteian, A., et al. (2024). Selective Aurora A-TPX2 Interaction Inhibitors Have In Vivo Efficacy as Targeted Antimitotic Agents. J Med Chem 67, 15521–15536. 10.1021/acs.jmedchem.4c01165.

5. Biswas, B., Huang, Y.H., Craik, D.J., and Wang, C.K. (2024). The prospect of substrate-based kinase inhibitors to improve target selectivity and overcome drug resistance. Chem Sci 15, 13130–13147. 10.1039/d4sc01088d.

6. Gimenez, D., Walko, M., Miles, J.A., Bayliss, R., Wright, M.H., and Wilson, A.J. (2024). Constrained TACC3 peptidomimetics for a non-canonical protein-protein interface elucidate allosteric communication in Aurora-A kinase. Chem Sci 16, 354–363. 10.1039/d4sc06100d.

7. Manschwetus, J.T., Bendzunas, G.N., Limaye, A.J., Knape, M.J., Herberg, F.W., and Kennedy, E.J. (2019). A Stapled Peptide Mimic of the Pseudosubstrate Inhibitor PKI Inhibits Protein Kinase A. Molecules 24. 10.3390/molecules24081567.

8. Crook, Z.R., Nairn, N.W., and Olson, J.M. (2020). Miniproteins as a Powerful Modality in Drug Development. Trends Biochem Sci 45, 332–346. 10.1016/j.tibs.2019.12.008.

9. Watson, J.L., Juergens, D., Bennett, N.R., Trippe, B.L., Yim, J., Eisenach, H.E., Ahern, W., Borst, A.J., Ragotte, R.J., Milles, L.F., et al. (2023). De novo design of protein structure and function with RFdiffusion. Nature 620, 1089–1100. 10.1038/s41586-023-06415-8.

10. Pacesa, M., Nickel, L., Schellhaas, C., Schmidt, J., Pyatova, E., Kissling, L., Barendse, P., Choudhury, J., Kapoor, S., Alcaraz-Serna, A., et al. (2025). One-shot design of functional protein binders with BindCraft. Nature 646, 483–492. 10.1038/s41586-025-09429-6.

11. Yang, W., Wang, S., Lee, G.R., Zhang, J.Z., Courbet, A., Juergens, D., Wang, X., Schlichthaerle, T., Abedi, M., Ragotte, R., et al. (2026). The past, present and future of de novo protein design. Nature 652, 1139–1152. 10.1038/s41586-026-10328-7.

12. Muratspahic, E., Feldman, D., Kim, D.E., Qu, X., Bratovianu, A.M., Rivera-Sanchez, P., Voss, J.H., Hertz, E.P.T., Jeppesen, M., Dimitri, F., et al. (2026). De novo design of miniproteins targeting GPCRs. Nature. 10.1038/s41586-026-10656-8.

13. Bavetsias, V., Faisal, A., Crumpler, S., Brown, N., Kosmopoulou, M., Joshi, A., Atrash, B., Perez-Fuertes, Y., Schmitt, J.A., Boxall, K.J., et al. (2013). Aurora isoform selectivity: design and synthesis of imidazo[4,5-b]pyridine derivatives as highly selective inhibitors of Aurora-A kinase in cells. J Med Chem 56, 9122–9135. 10.1021/jm401115g.

14. Rennie, Y.K., McIntyre, P.J., Akindele, T., Bayliss, R., and Jamieson, A.G. (2016). A TPX2 Proteomimetic Has Enhanced Affinity for Aurora-A Due to Hydrocarbon Stapling of a Helix. ACS Chem Biol 11, 3383–3390. 10.1021/acschembio.6b00727.

15. Burgess, S.G., Oleksy, A., Cavazza, T., Richards, M.W., Vernos, I., Matthews, D., and Bayliss, R. (2016). Allosteric inhibition of Aurora-A kinase by a synthetic vNAR domain. Open Biol 6. 10.1098/rsob.160089.

16. Dawber, R.S., Gimenez, D., Batchelor, M., Miles, J.A., Wright, M.H., Bayliss, R., and Wilson, A.J. (2024). Inhibition of Aurora-A/N-Myc Protein-Protein Interaction Using Peptidomimetics: Understanding the Role of Peptide Cyclization. Chembiochem 25, e202300649. 10.1002/cbic.202300649.

17. Boi, D., Fianco, G., Polverino, F., Fiorentino, F., Mastrangelo, A., Rossi, S., Rubini, E., Rosignoli, S., Troilo, F., Antonelli, M.R., et al. (2026). The ATC12 small molecule inhibits the Aurora-A/TPX2 interaction and impairs the proliferation of breast cancer cells. Cell Death Dis 17. 10.1038/s41419-026-08579-3.

18. Bayliss, R., Sardon, T., Vernos, I., and Conti, E. (2003). Structural basis of Aurora-A activation by TPX2 at the mitotic spindle. Mol Cell 12, 851–862. 10.1016/s1097-2765(03)00392-7.

19. Burgess, S.G., Mukherjee, M., Sabir, S., Joseph, N., Gutierrez-Caballero, C., Richards, M.W., Huguenin-Dezot, N., Chin, J.W., Kennedy, E.J., Pfuhl, M., et al. (2018). Mitotic spindle association of TACC3 requires Aurora-A-dependent stabilization of a cryptic alpha-helix. EMBO J 37. 10.15252/embj.201797902.

20. Holder, J., Miles, J.A., Batchelor, M., Popple, H., Walko, M., Yeung, W., Kannan, N., Wilson, A.J., Bayliss, R., and Gergely, F. (2024). CEP192 localises mitotic Aurora-A activity by priming its interaction with TPX2. EMBO J 43, 5381–5420. 10.1038/s44318-024-00240-z.

21. Park, J.G., Jeon, H., Shin, S., Song, C., Lee, H., Kim, N.K., Kim, E.E., Hwang, K.Y., Lee, B.J., and Lee, I.G. (2023). Structural basis for CEP192-mediated regulation of centrosomal AURKA. Sci Adv 9, eadf8582. 10.1126/sciadv.adf8582.

22. Richards, M.W., Burgess, S.G., Poon, E., Carstensen, A., Eilers, M., Chesler, L., and Bayliss, R. (2016). Structural basis of N-Myc binding by Aurora-A and its destabilization by kinase inhibitors. Proc Natl Acad Sci U S A 113, 13726–13731. 10.1073/pnas.1610626113.

23. Zorba, A., Buosi, V., Kutter, S., Kern, N., Pontiggia, F., Cho, Y.J., and Kern, D. (2014). Molecular mechanism of Aurora A kinase autophosphorylation and its allosteric activation by TPX2. Elife 3, e02667. 10.7554/eLife.02667.

24. Zorba, A., Nguyen, V., Koide, A., Hoemberger, M., Zheng, Y., Kutter, S., Kim, C., Koide, S., and Kern, D. (2019). Allosteric modulation of a human protein kinase with monobodies. Proc Natl Acad Sci U S A 116, 13937–13942. 10.1073/pnas.1906024116.

25. Bennett, N.R., Coventry, B., Goreshnik, I., Huang, B., Allen, A., Vafeados, D., Peng, Y.P., Dauparas, J., Baek, M., Stewart, L., et al. (2023). Improving de novo protein binder design with deep learning. Nat Commun 14, 2625. 10.1038/s41467-023-38328-5.

26. Burgess, S.G., and Bayliss, R. (2015). The structure of C290A:C393A Aurora A provides structural insights into kinase regulation. Acta Crystallogr F Struct Biol Commun 71, 315–319. 10.1107/S2053230X15002290.

27. Rejnowicz, E., Batchelor, M., Leen, E., Ahangar, M.S., Burgess, S.G., Richards, M.W., Kalverda, A.P., and Bayliss, R. (2024). Exploring the dynamics and interactions of the N-myc transactivation domain through solution nuclear magnetic resonance spectroscopy. Biochem J 481, 1535–1556. 10.1042/BCJ20240248.

28. Willems, E., Dedobbeleer, M., Digregorio, M., Lombard, A., Lumapat, P.N., and Rogister, B. (2018). The functional diversity of Aurora kinases: a comprehensive review. Cell Div 13, 7. 10.1186/s13008-018-0040-6.

29. Abdul Azeez, K.R., Chatterjee, S., Yu, C., Golub, T.R., Sobott, F., and Elkins, J.M. (2019). Structural mechanism of synergistic activation of Aurora kinase B/C by phosphorylated INCENP. Nat Commun 10, 3166. 10.1038/s41467-019-11085-0.

30. Huse, M., and Kuriyan, J. (2002). The conformational plasticity of protein kinases. Cell 109, 275–282. 10.1016/s0092-8674(02)00741-9.

31. Attwood, M.M., Fabbro, D., Sokolov, A.V., Knapp, S., and Schioth, H.B. (2021). Trends in kinase drug discovery: targets, indications and inhibitor design. Nat Rev Drug Discov 20, 839–861. 10.1038/s41573-021-00252-y.

32. Johnson, J.L., Yaron, T.M., Huntsman, E.M., Kerelsky, A., Song, J., Regev, A., Lin, T.Y., Liberatore, K., Cizin, D.M., Cohen, B.M., et al. (2023). An atlas of substrate specificities for the human serine/threonine kinome. Nature 613, 759–766. 10.1038/s41586-022-05575-3.

33. Dunbrack, R.L. Rēs ipSAE loquuntur: What’s wrong with AlphaFold’s ipTM score and how to fix it. BioRxiv. 10.1101/2025.02.10.637595.

34. Damodaran, A.P., Gavard, O., Gagne, J.P., Rogalska, M.E., Behera, A.K., Mancini, E., Bertolin, G., Courtheoux, T., Kumari, B., Cailloce, J., et al. (2025). Proteomic study identifies Aurora-A-mediated regulation of alternative splicing through multiple splicing factors. J Biol Chem 301, 108000. 10.1016/j.jbc.2024.108000.

35. Lu, R.M., Hsu, H.E., Perez, S., Kumari, M., Chen, G.H., Hong, M.H., Lin, Y.S., Liu, C.H., Ko, S.H., Concio, C.A.P., et al. (2024). Current landscape of mRNA technologies and delivery systems for new modality therapeutics. J Biomed Sci 31, 89. 10.1186/s12929-024-01080-z.

36. Garralda, E., Beaulieu, M.E., Moreno, V., Casacuberta-Serra, S., Martinez-Martin, S., Foradada, L., Alonso, G., Masso-Valles, D., Lopez-Estevez, S., Jauset, T., et al. (2024). MYC targeting by OMO-103 in solid tumors: a phase 1 trial. Nat Med 30, 762–771. 10.1038/s41591-024-02805-1.

37. Dauparas, J., Anishchenko, I., Bennett, N., Bai, H., Ragotte, R.J., Milles, L.F., Wicky, B.I.M., Courbet, A., de Haas, R.J., Bethel, N., et al. (2022). Robust deep learning-based protein sequence design using ProteinMPNN. Science 378, 49–56. 10.1126/science.add2187.

38. Winter, G., Lobley, C.M., and Prince, S.M. (2013). Decision making in xia2. Acta Crystallogr D Biol Crystallogr 69, 1260–1273. 10.1107/S0907444913015308.

39. McCoy, A.J., Grosse-Kunstleve, R.W., Adams, P.D., Winn, M.D., Storoni, L.C., and Read, R.J. (2007). Phaser crystallographic software. J Appl Crystallogr 40, 658–674. 10.1107/S0021889807021206.

40. Jumper, J., Evans, R., Pritzel, A., Green, T., Figurnov, M., Ronneberger, O., Tunyasuvunakool, K., Bates, R., Zidek, A., Potapenko, A., et al. (2021). Highly accurate protein structure prediction with AlphaFold. Nature 596, 583–589. 10.1038/s41586-021-03819-2.

41. Emsley, P., and Cowtan, K. (2004). Coot: model-building tools for molecular graphics. Acta Crystallogr D Biol Crystallogr 60, 2126–2132. 10.1107/S0907444904019158.

42. Liebschner, D., Afonine, P.V., Baker, M.L., Bunkoczi, G., Chen, V.B., Croll, T.I., Hintze, B., Hung, L.W., Jain, S., McCoy, A.J., et al. (2019). Macromolecular structure determination using X-rays, neutrons and electrons: recent developments in Phenix. Acta Crystallogr D Struct Biol 75, 861–877. 10.1107/S2059798319011471.

43. Williams, C.J., Headd, J.J., Moriarty, N.W., Prisant, M.G., Videau, L.L., Deis, L.N., Verma, V., Keedy, D.A., Hintze, B.J., Chen, V.B., et al. (2018). MolProbity: More and better reference data for improved all-atom structure validation. Protein Sci 27, 293–315. 10.1002/pro.3330.

44. Joosten, R.P., Joosten, K., Cohen, S.X., Vriend, G., and Perrakis, A. (2011). Automatic rebuilding and optimization of crystallographic structures in the Protein Data Bank. Bioinformatics 27, 3392–3398. 10.1093/bioinformatics/btr590.

45. Krissinel, E., and Henrick, K. (2007). Inference of macromolecular assemblies from crystalline state. J Mol Biol 372, 774–797. 10.1016/j.jmb.2007.05.022.

46. Holm, L., Laiho, A., Toronen, P., and Salgado, M. (2023). DALI shines a light on remote homologs: One hundred discoveries. Protein Sci 32, e4519. 10.1002/pro.4519.

47. Demichev, V., Messner, C.B., Vernardis, S.I., Lilley, K.S., and Ralser, M. (2020). DIA-NN: neural networks and interference correction enable deep proteome coverage in high throughput. Nat Methods 17, 41–44. 10.1038/s41592-019-0638-x.

48. Katoh, K., Rozewicki, J., and Yamada, K.D. (2019). MAFFT online service: multiple sequence alignment, interactive sequence choice and visualization. Brief Bioinform 20, 1160–1166. 10.1093/bib/bbx108.

49. Crooks, G.E., Hon, G., Chandonia, J.M., and Brenner, S.E. (2004). WebLogo: a sequence logo generator. Genome Res 14, 1188–1190. 10.1101/gr.849004.

