## Supplementary Information for "Selective miniprotein inhibitors of Aurora-A kinase designed using interaction-motif scaffolding"

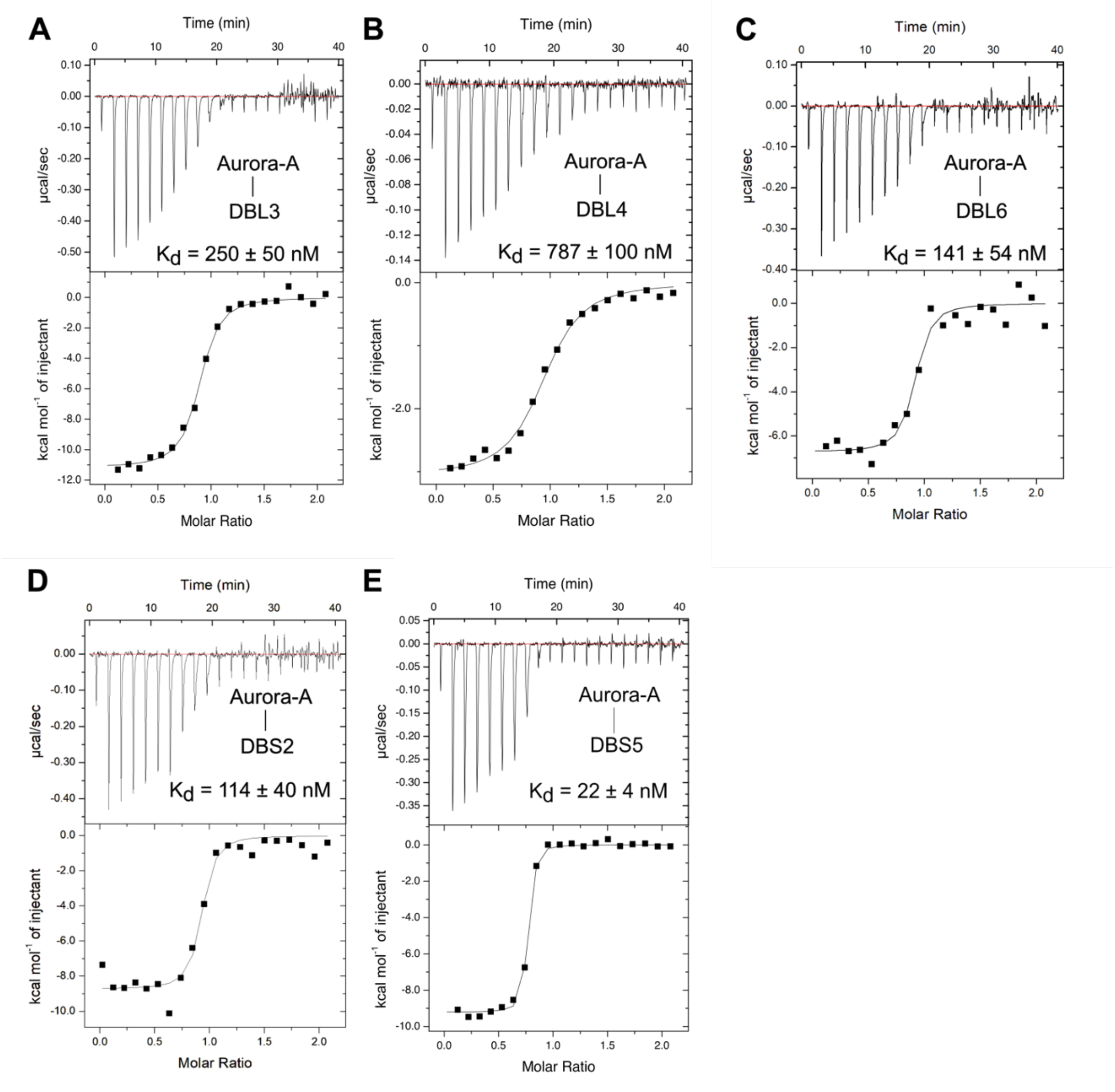


**Figure S1: ITC characterisation of designed miniproteins to Aurora-A kinase.**

**A-E:** Isothermal titration calorimetry experiments showing titration of Aurora-A kinase domain (122–403 C290A C393A D274N) into (A) DBL3, (B) DBL4, (C) DBL6, (D) DBS2, (E) DBS5. The measured $K_d$ and fitted n values are from two experimental repeats.

**A**

```
               ! !!  !  !
N-myc       PPSWVTEMLLENELWG

DBS1        PPEWVIQLLKEILEEV
DBS5        LPEWVYQLLLELLTTS
DBL3        PPDWVLEILDEIIKNG
DBL5        LPEWVIKIFEEFKAKG
DBL6        LPEWVDKIIEEIVENG
DBL2        LPDWVIKIAEEMVTKG
DBL4        LPDWVYAIIDEALENG
DBS2        PPDWVYQLIIEVLEQE
DBL1        PPQWVLDILAEAKAKG
DBL8        PPDWVYKILEEAKEKG
DBL10       LPDWVVKLVEEFRKNG
DBS3        PPEWVIKLLKELIEEL
DBS4        PPDWVIELLAEVLGVS
DBI1        PPDWVTELIVENALWG
DBI2        PPDWVTELFEENKKWG
              *:**   :  *
```

**B** **C**

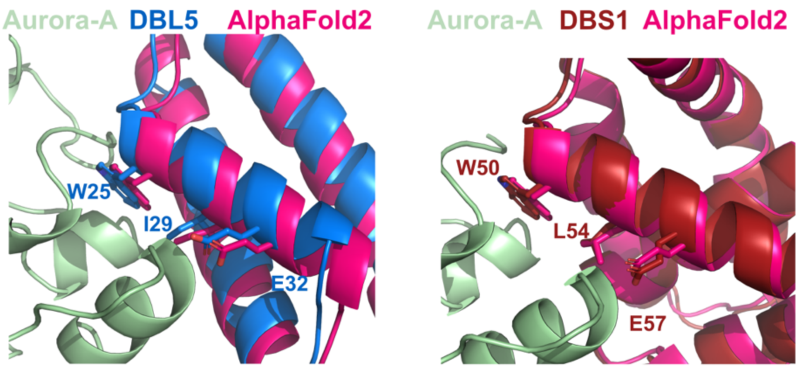


**D**

| Region of Aurora-A | N-myc 61-89 | N-myc 74-89 Templated helix | DBL5 | DBS1 |
|---|---|---|---|---|
| Gly-rich loop N-lobe | K143 | | | K143 |
| αB-αC helices N-lobe | A172, G173, H176, R180, E183 | | A172, G173, V174 | E170, K171, A172 |
| Activation loop C-lobe | W277, V279, H280, P282, R286, T288, L289, A290, G291, L293, L296, I301 | W277, L289, A290, G291, L293, L296, I301 | W277, T287, T288, L289, A290, G291, L293, L296, I301, E302, G303, R304 | W277, T288, L289, A290, G291, L293, L296, M300, I301, E302, G303 |
| αG helix C-lobe | T333, Y334, Q335, Y338 | T333, Y334, Q335, Y338 | T333, Y334, Q335, Y338 | T333, Y334, Q335, Y338 |

**Figure S2: Comparison of Aurora-A binding regions between N-myc and miniproteins**

**A:** Alignment of designed binder central helix sequences, based on N-myc scaffolded sequence.

**B:** Superposition of DBL5/Aurora-A crystal structure and AlphaFold2 models.

**C:** Superposition of DBS1/Aurora-A crystal structure and AlphaFold2 models.

**D:** Lists of Aurora-A regions and residues at the interface with N-Myc, DBL5 or DBS1.

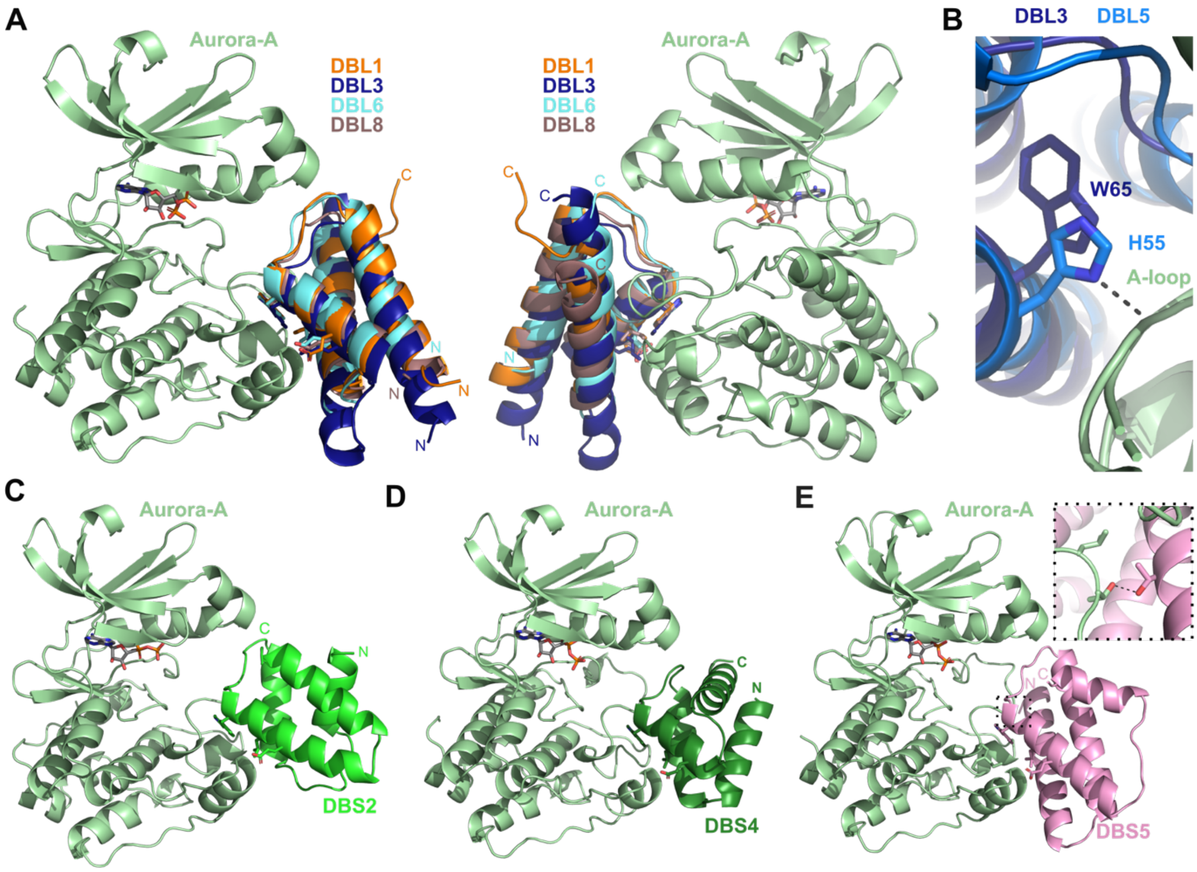


**Figure S3: Crystal structures of further binder/Aurora-A complexes**

**A:** Overlay of the structures of 4 Aurora-A/binder complexes, all involving miniproteins based on 3-helix bundles. DBL1 is shown in orange, with DBL3 in dark blue, DBL6 in cyan and DBL8 in grey-pink.
**B:** Magnified view of interaction between DBL5 H55 and A-loop. Note equivalent residue in DBL3 is W65, which does not form an H-bond with the A-loop.
**C:** Crystal structure of Aurora-A/DBS2 complex.
**D:** Crystal structure of Aurora-A/DBS4 complex.
**E:** Crystal structure of Aurora-A/DBS5 complex. The inset shows the interaction between Thr10 of DBS5 and Thr288 of the Aurora-A activation loop.

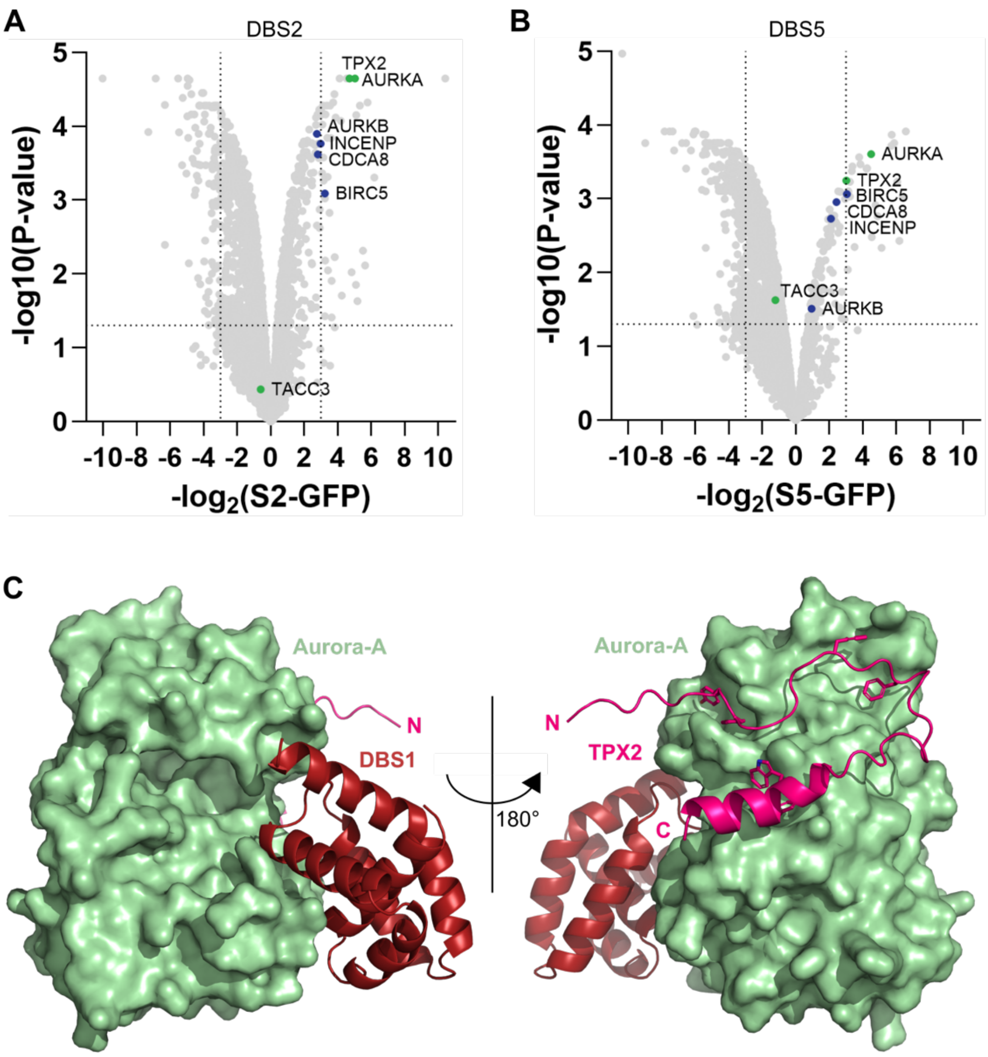


**Figure S4: Designed binders interact with the Aurora-A/TPX2 complex**

**A-B:** Volcano plot analysis showing log2 fold-change and significance (-log10 p-value) comparing protein enrichment with GFP-tagged designed binders (A) DBS2 or (B) DBS5 against GFP-only control. HeLa cells were transfected, as in Figure 6A, and 100 ng/ml nocodazole was added during the final 20 hours of incubation to arrest cells in mitosis. GFP-immunoprecipitation was performed and samples analysed by LC-MS.
**C**: AlphaFold3 model of a ternary complex between Aurora-A (light green), DBS1 (dark red) and TPX2 (pink). There does not appear to be any clash between DBS1 and TPX2 on the surface of Aurora-A.

### Supplementary Table S1

### Crystallographic data

| | Aurora-A:DBL1 | Aurora-A:DBL3 | Aurora-A:DBL5 | Aurora-A:DBL6 | Aurora-A:DBL8 | Aurora-A:DBS1 | Aurora-A:DBS2 | Aurora-A:DBS4 | Aurora-A:DBS5 |
|---|---|---|---|---|---|---|---|---|---|
| X-ray source | DLS i04 | DLS i03 | DLS i04 | DLS i04 | DLS i04 | DLS i04 | DLS i03 | DLS i04 | DLS i03 |
| Wavelength (Å) | 0.9537 | 0.9763 | 0.9537 | 0.9537 | 0.9537 | 0.9537 | 0.9762 | 0.9537 | 0.9763 |
| Resolution range (Å) | 63.52 – 2.36 | 47.49 – 2.07 | 54.57- 1.79 | 57.22 – 2.35 | 55.83 – 2.37 | 51.27 – 2.00 | 58.96 – 2.90 | 60.45 – 2.23 | 71.67 – 2.28 |
| Space group | P21 21 21 | P 21 21 21 | P 21 21 21 | P 21 21 21 | P 21 21 21 | P 21 21 21 | P 32 2 1 | P 41 21 2 | P 31 2 1 |
| Unit cell (a,b,c Å) (α,β,γ, °) | 49.77 86.03 94.18 90.00 90.00 90.00 | 40.27 91.32 111.19 90.00 90.00 90.00 | 56.77 68.85 89.23 90.00 90.00 90.00 | 54.73 75.48 87.74 90.00 90.00 90.00 | 57.39 71.76 88.88 90.00 90.00 90.00 | 64.41 77.42 84.24 90.00 90.00 90.00 | 84.05 84.05 201.1 90.00 90.00 120.00 | 88.33 88.33 240.29 90.00 90.00 90.00 | 87.09 87.09 215.26 90.00 90.00 120.00 |
| Unique reflections | 13349 | 25854 | 33504 | 12251 | 11954 | 28883 | 18955 | 47348 | 41554 |
| Multiplicity | 9.3 | 13.3 | 13.6 | 8.3 | 9.5 | 13.6 | 16.6 | 26.4 | 14.6 |
| Completeness (%) | 78.3* | 100.0 | 99.3 | 78.5* | 78.9* | 99.2 | 99.9 | 99.9 | 94.3 |
| Mean I/σ(I) | 20.7 | 14.5 | 10.7 | 28.2 | 19.3 | 13.8 | 29.6 | 14.1 | 14.6 |
| $CC_{1/2}$ | 1.0 | 1.0 | 1.0 | 1.0 | 1.0 | 1.0 | 1.0 | 1.0 | 1.0 |
| Refinement statistics | | | | | | | | | |
| $R_{work}$ | 0.201 | 0.227 | 0.195 | 0.197 | 0.187 | 0.187 | 0.204 | 0.201 | 0.19 |
| $R_{free}$ | 0.250 | 0.267 | 0.238 | 0.259 | 0.260 | 0.228 | 0.244 | 0.249 | 0.231 |
| Number of non- hydrogen atoms | 2673 | 2739 | 2907 | 2638 | 2551 | 3044 | 5201 | 5442 | 5914 |
| Macromolecule | 2577 | 2664 | 2611 | 2525 | 2471 | 2933 | 5090 | 5054 | 5605 |
| Ligand | 28 | 27 | 63 | 28 | 27 | 27 | 58 | 87 | 68 |
| Water | 68 | 48 | 232 | 85 | 53 | 84 | 53 | 301 | 241 |
| Protein residues | 325 | 330 | 324 | 316 | 313 | 359 | 660 | 646 | 706 |
| RMS bonds | 0.45 | 0.46 | 0.56 | 0.45 | 0.47 | 0.48 | 0.48 | 0.51 | 0.50 |
| RMS angles | 0.82 | 0.81 | 1.17 | 0.86 | 0.89 | 0.90 | 0.88 | 0.95 | 0.97 |
| Ramachandran favoured (%) | 97 | 97 | 96 | 98 | 97 | 97 | 97 | 97 | 97 |
| MolProbity Clashscore | 2.0 | 1.0 | 8.0 | 3.0 | 2.0 | 2.0 | 2.0 | 3.0 | 2.0 |
| Average B-factor | 21.0 | 68.0 | 39.0 | 38.0 | 44.0 | 57.0 | 54.0 | 63.0 | 70.0 |
| PDB ID code | 9S0K | 9S0W | 9RVK | 9S1G | 9S61 | 9S14 | 9SQ7 | 9SV0 | 9S7T |

*STARANISO data
